# Longitudinal study of liver disease progression in the PEX1-Gly844Asp mouse model of mild Zellweger Spectrum Disorder

**DOI:** 10.1101/2025.05.08.652960

**Authors:** Lingxiao Chen, Hong Choi, Catherine Argyriou, Monica Hsieh, Erminia Di Pietro, Wei Cui, Esther Nuebel, Caroline Daneault, Matthieu Ruiz, Daniel Charpentier, David Rhainds, Joseph G Hacia, Van-Hung Nguyen, Zu-Hua Gao, Nancy Braverman

## Abstract

**Introduction:** Zellweger spectrum disorder (ZSD) is an autosomal recessive disorder caused by mutations in any of 13 *PEX* genes encoding proteins required for peroxisome assembly and function. Chronic liver disease is one of the major clinical manifestations in patients and impacts quality of life and survival. However, the pathophysiology of liver disease is ZSD remains largely unknown, and current interventions are limited. To further study the liver disease mechanism, we use the PEX1-Gly844Asp (G844D) mouse model for mild ZSD, which was previously shown to develop hepatomegaly and cholestasis, similar to ZSD patients.

**Methods:** The natural history of hepatopathy was broadly characterized in PEX1-G844D mice and littermate controls from 1 to 18 months of age using liver histology, electron microscopy, cultured hepatocytes and blood. Metabolite and mechanism analysis included liver functions, respiratory chain dynamics, lipidomics, peroxisome metabolites, gene and protein expression assays.

**Results:** PEX1-G844D mice featured liver disease progression from hepatomegaly (1 month) to cluster cell death (4 months), hepatosteatosis (6 months), inflammation (8 months), fibrosis, and hepatic cancer (12 and 15 months). Hepatocyte proliferation and reduced glycogen was observed across all ages. Measurement of peroxisomal functions showed defective peroxisomal import and secondary mitochondrial defects in cultured hepatocytes. In blood and liver, plasmalogens were decreased, and C26:0 lyso-phosphatidylcholine and C27 bile acid intermediates were elevated. In liver, we observed accumulation of triglycerides and cholesterol, and reduced membrane phospholipids and sphingolipids. In contrast, in serum we observed reduced triglycerides, cholesterol and membrane lipids. Gene expression profiles confirmed by immunoblotting supported reduced hepatic *de novo* lipogenesis, increased hepatic lipid uptake and oxidation, PPARα activation, and modulated glucose and glycogen metabolism. Liver X receptor agonist (T0901317) applied to cultured hepatocytes enhanced hepatic lipogenesis and lipid secretion, but aggravated steatosis.

**Conclusion:** Taken together, these results suggested the following mechanisms of hepatopathy progression. We propose that global peroxisome dysfunction (1) causes PPARα activation, leading to chronic hyperplasia and partially contributing to disrupted hepatic lipid homeostasis with hepatosteatosis, and (2) underlies chronic hypoglycemia, causing hypoinsulinemia and contributing to reduced hepatic lipogenesis and systemic lipid deficiency. Growth restriction in the mouse model and in ZSD patients could be attributable to systemic lipid deficiency. Our mechanistic delineation of the pathophysiology provides other additional novel potential therapeutic targets to halt liver disease in ZSD.

## INTRODUCTION

Peroxisomes are spherical, ubiquitous, single membrane-bound organelles numbering up to several hundred in mammalian cells. Each peroxisome contains more than 50 enzymes required for several vital metabolic pathways, which include β-oxidation of very long chain fatty acids (VLCFAs), α-oxidation of methyl-branched chain phytanic acids, biosynthesis of bile acids and ether phospholipids (plasmalogens), and metabolism of hydrogen peroxide. *PEX* genes encode peroxin proteins essential for peroxisome biogenesis, which includes membrane assembly, peroxisome division and matrix enzyme import. Biallelic mutations in any of 13 *PEX* genes will cause Zellweger Spectrum Disorder (ZSD), a heterogenous disorder resulting in multi-systemic complications affecting brain, hearing, vision, adrenals, bone and liver (1).

Mutations in the *PEX1* gene are the most common cause of ZSD, representing roughly 70% of ZSD patients in North America (2–4). PEX1 and PEX6 forms the peroxisome ‘exportomer complex (5) to export and recycle the peroxisome enzyme receptors back to cytosol for additional rounds of peroxisomal enzyme import. Defective PEX1-PEX6 export hence impairs peroxisome function by disrupting peroxisome matrix enzyme import (6, 7). *PEX1*-c.[2528G>A] accounts for roughly 30% of ZSD-causing alleles, and encodes the missense protein PEX1-p.[Gly843Asp] (or G843D) with residual function (8, 9). Patients with at least one PEX1-G843D allele exhibit a milder clinical phenotype with progressive multi-systemic disorder due to life-long peroxisome dysfunction (10, 11).

Liver disease is a major symptom of ZSD. In general, infants with severe ZSD experienced transient neonatal jaundice, cholestasis with conjugated hyperbilirubinemia at early infancy, hepatomegaly and some were reported to develop fibrosis, cirrhosis, and rarely, liver failure within the first year of life (12, 13). In patients with intermediate-mild ZSD, transient neonatal jaundice, hepatomegaly, elevated plasma levels of liver transaminases and coagulopathy have been reported (14, 15). Although clinical manifestations of liver disease are generally absent in milder ZSD phenotypes, a subset of patients progress to fibrosis, cirrhosis and portal hypertension, which causes esophageal varices, gastrointestinal bleeding episodes and ascites. In a few cases, hepatocellular carcinoma resulting in death was reported (16) or dysplastic tumors were found on autopsy (14, 15).

To investigate the course and mechanism of disease progression in ZSD, and to identify clinical endpoints for future therapeutic studies, we engineered a mouse model of mild ZSD, which carries the PEX1-G843D equivalent knock-in mutation (PEX1-G844D). Preliminary liver studies showed that this mouse model recapitulates many of human disease characteristics, including growth retardation, hepatomegaly, cholestasis, abnormal peroxisome metabolites and liver functions (17). Other research groups using independently derived PEX1-G844D murine models reported liver hyperplasia, steatosis, fibrosis, mitochondrial dysfunction and abnormal carbohydrate metabolism up to 6 months of age (18, 19). In this study, we extend and detail the natural history of liver disease development in PEX1-G844D mice up to 18 months of age, and identify the disrupted lipid metabolic pathways underlying liver disease progression and dyslipidemia. We link the disrupted lipid homeostasis with defects in glucose metabolism and mitochondrial dysfunction. Finally, we propose preclinical endpoints for future preclinical studies for therapeutic interventions.

## MATERIALS AND METHODS

### Animal husbandry

PEX1-G844D mice were maintained on a mixed 129/SvEv and C57BL/6N Taconic background and showed a stable 70% 129/SvEv and 30% C57BL/6N Taconic as reported (20). Peroxisome metabolites, microarray analysis, untargeted lipidomic analysis and histology were conducted on liver tissues. Afterwards, all experiments were performed on PEX1-G844D homozygotes (F1) derived from crossing PEX1-G844D heterozygotes congenic on 129/SvEv to PEX1-G844D heterozygotes congenic on C57BL/6N background (21). F1 mice were therefore always 50% 129/SvEv and 50% C57BL/6N and showed 100% survival. Mice were housed at the RI-MUHC Glen site animal care facility with *ad libitum* access to food and water. All experiments were performed at the RI-MUHC Glen site and were approved by the Research Institute of the McGill University Health Centre Animal Care Committee. Euthanasia was performed by CO2 asphyxiation under isoflurane anaesthesia. Both males and females were used for all experiments. Wild-type and PEX1-G844D heterozygous (wt/G844D) mice were used as littermate controls.

### Genotyping

For routine genotyping, genomic DNA was isolated from ear punches of 21-day old mice by incubating in 75μl alkaline lysis buffer (25mM NaOH, 0.2mM Na2EDTA) at 95°C for 20 min, then neutralized with 250μl 40mM Tris-HCl. Genotypes were determined by PCR amplification (forward primer 5’-TCAATGTGTCCAGCACCTTC-3’; reverse primer 5’-TATGGAACGGAATGAGGC-3’) that produces amplicons of different sizes depending on the presence of a 177 bp residual Neo cassette fragment in intron 13 near the PEX1 c.2531G>A mutation in exon 15. The *Pex1-*c.2531G>A (PEX1-p.Gly844Asp) allele yields a 852 base pair product and the wild-type allele yields a 672 base pair product (17, 22).

### Microarray analysis

Total RNA was isolated from mouse liver dissected at 124 days of age and subjected to global gene expression analysis on GeneChip® Mouse Genome 430a2.0 Arrays (Affymetrix, Santa Clara CA, USA) designed to interrogate over 14,000 transcripts. As previously described (23), the resulting .CEL files were preprocessed using the WebArray software (24) that uses the RMA (Robust Multi-array Average) algorithm (25) to generate log2-scaled expression values for each transcript. Using the LIMMA (Linear Models for Microarray) package (26), we selected probe sets showing absolute fold change greater than 1.2, and a false discovery rate (FDR) value less than 0.05. The FDR values were calculated by adjusting raw P-values using the SPLOSH (Spacing LOESS Histogram) method (27).

### Quantitative Real-Time PCR (qPCR)

Total RNA was extracted from (∼50 mg) liver using TRIZOL™ (Invitrogen, 15596026, Carlsbad, CA). For reverse transcription and cDNA amplification we used: iScript® cDNA Synthesis Supermix reverse transcriptase (Bio-Rad, 1708891), qRT-PCR LunaScript RT SuperMix Kit (NEB, E3010) and CFX96 Touch Real-Time PCR Detection System (Bio-Rad). Gene expression was normalized to murine beta-2-microglobulin (*B2m*) and beta-glucuronidase (*Gusb*) using Bio-Rad CFX Maestro software. All qPCR experiments were performed twice per sample. In each experiment, we used 3 biological and 2 technical replicates per sample. Primer sets used for qPCR experiments are listed in **Supplemental Table 1**.

### Immunoblotting

50mg flash-frozen mouse livers were homogenized in RIPA lysis buffer. Lysates (30 μg) were separated on 10% polyacrylamide gel and transferred to nitrocellulose membrane. Membranes were blocked and hybridized in 5% milk with primary antibodies: 1:500 mouse anti-human PEX1 (BD Bioscience, 611719), 1:2000 rabbit anti-human PEX5 (ProteinTech 12545-1-AP), 1:2000 rabbit anti-human PEX6 (gift from Dr. Gabriele Dodt, University of Tübingen),1:1000 rabbit anti-mouse APOB (ProteinTech, 20578-1-AP) and VLDLR (ProteinTech, 19493-1-AP), 1:1500 rabbit anti-mouse PKLR (ProteinTech, 22456-1-AP), 1:5000 rabbit anti-mouse HADHA (Abcam, ab54477), SOD2 (Abcam, ab13533), 1:5000 rabbit anti-mouse G6PC (ProteinTech, 22169-1-AP), 1:1000 rabbit anti-mouse FASN (Cell Signalling, C2065), 1:5000 mouse anti-human MTP (BD, 612022), 1:1000 mouse anti-human SREBP-1 (Novus, NB600-582), 1:1000 rabbit anti-mouse CD36 (Novus, NB400-144), 1:17000 rabbit anti-human β-tubulin (Abcam, ab6046), followed by appropriate HRP-conjugated secondary antibodies, and visualized by ECL using an Amersham 600 platform. Band quantification (densitometry) was done using ImageJ (NIH).

### Liver histology, immunohistochemistry and immunofluorescence

#### Histology

5μm paraffin-embedded sections were stained by hematoxylin and eosin (H&E) (Sigma Aldrich, GHS116, HT110116), periodic-acid Schiff (PAS) (Sigma Aldrich, 395B-1KT) and Sirius red (Abcam, ab150681). For Oil Red O stain, 10μm cryo-sections were stained with 60% Oil Red O solution prepared from powder (VWR, 0684). Slides were visualized using an Olympus BX51 microscope at 10x and 40x magnification; images were captured using an Olympus CCD camera and MagnaFire software (Olympus). Staining area and intensity were determined using ImageJ (NIH). 3 fields of view per section (animal) were measured from 3 animals per genotype from each age group.

#### Immunohistochemistry

5μm paraffin-embedded liver sections were antigen retrieved and blocked. Sections were incubated in primary antibody overnight at 4°C. After washing, secondary antibody was applied for 1 hour incubation, followed by VECTASTAIN Elite ABC kit (Vector Laboratories, PK6101), DAB immunostaining (Abcam, ab64238), and hematoxylin counterstaining (Sigma Aldrich). Primary antibodies used were 1:200 rabbit anti-mouse SOD2 (Abcam, ab13533) and Glypican-3 (Abcam ab66596), 1:100 mouse anti-human Ki67 (Abcam, ab16667), 1:150 mouse anti-human glutamate synthase (Millipore Sigma, MAB302). Ki67 index was determined by manually labeling Ki67 positive nucleus using ImageJ to count the number. 3 fields of view per section (animal) was counted from 3 animals per genotype from each age group.

#### Immunofluorescence

5μm paraffin-embedded liver sections were antigen retrieved and blocked. Sections were incubated in primary antibody overnight at 4°C, followed by secondary antibody with fluorescence labeling for 1 hour. Autofluorescence was quenched with 0.3% Sudan black B in 70% ethanol, followed by mounting with DAPI (Invitrogen, P36935). Images were visualized by Leica DMI6000B microscope with DFC345FX camera and 106 LASX software (Richmond Hill, Canada). Primary antibodies used were 1:300 rabbit anti-human PEX14, SKL (gift from Steven Gould, John Hopkins University), 1:200 rabbit anti-human catalase (Aoxre 23416), 1:150 mouse anti-human PMP70 (Sigma, SAB4200181); secondary antibodies: 1:400 anti-rabbit 488 (Invitrogen, A21206), 1:300 anti-mouse 594 (Invitrogen, A11005).

### Transmission electron microscopy

Protocol was adapted from *Fahimi, 2017* (28). Following euthanasia, mouse liver was cannulated and perfused with 0.15M sodium cacodylate buffer (pH 7.2) containing 0.05M calcium chloride and 1.25% glutaraldehyde for 10 min through inferior vena cava. Liver was removed and immersed in 0.1M sodium cacodylate buffer (pH 7.2) containing 0.01M calcium chloride, 1.25% glutaraldehyde and 4% paraformaldehyde for fixation overnight. Liver tissue was washed, sliced at 50μm with vibratome, and stained by 3,3′-Diaminobenzidine tetrahydrochloride medium (0.2% DAB, 0.15% hydrogen peroxide in 0.01M Teorell-Stenhagen buffer, pH 10.5) for 1 hr at room temperature. Liver sections were then post fixed with 1% osmium tetroxide, washed, and processed by 2% uranyl acetate in 0.05M Na-H Maleate buffer for 30 min at 4°C. Samples were dehydrated with increasing concentrations of ethanol, infiltrated with increasing concentrations of Epon, then polymerized and embedded in pure Epon at 60°C for 48 hrs. 100 nm sections were cut and placed onto a 200 mesh copper grid, counterstained with uranyl acetate and alkaline lead citrate for 2 min, and imaged with the FEI Tecnai 12 BioTwin TEM equipped with the AMT XR80C CCD camera at an accelerating voltage of 120 kV.

Mitochondria size was determined by manually tracing around each individual mitochondrion and using ImageJ (NIH) to calculate its area. 5–14 mitochondria were measured per animal for a total of 46 mutant and 30 control mitochondria scored from 4 mice of each genotype. Peroxisome number was calculated manually and normalized by area using ImageJ. 1-3 fields of view were measured per animal for total of 10 mutant and 9 control images analyzed from 5 mice of each genotype.

### Biochemical analyses

Flash frozen liver tissue from PEX1-G844D mutants and wild-type littermates at age 1-9 months were analysed for VLCFAs, phytanic and pristanic acids, and bile acids using gas chromatography-mass spectrometry (GC-MS/MS) as previously reported (29, 30). C26:0-lysophosphatidylcholine and plasmalogens were measured in dried bloodspots from 2-month-old mice on filter paper (Whatman 903, GE LifeScience) using liquid chromatography-mass spectrometry (LC-MS/MS) as previously described (17). Serum was analyzed at Comparative Medicine Animal Resource Centre laboratory (McGill University) for ALT, ALP, AST, Glucose, cholesterol and triglyceride with enzymatic colorimetric tests using the SYNCHRON® system. Serum levels of alpha-fetoprotein (AFP) and insulin were measured by solid phase sandwich ELISA: Mouse AFP Quantikine ELISA Kit (R&D system, MAFP00) and Mouse Ultrasensitive Insulin ELISA kit (ALPCO, #80-INSMSU-E01). 10uL and 25uL of serum or standard was loaded in duplicates for AFP and Insulin immunoassay, respectively, and performed as outlined in product instruction manual. Colorimetric assay was read at wavelength 450nm by BioTek Epoch microplate reader immediately after reaction.

### Untargeted lipidomics analysis and lipoprotein precipitation

#### Untargeted lipidomics

Lipid extraction, sample and data analysis were performed using a previously validated untargeted lipidomic workflow (31–33). Briefly, lipids were extracted from 100uL of serum and spiked with six internal standards: LPC 13:0, PC19:0/19:0, PC14:0/14:0, PS12:0/12:0, PG15:0/15:0, and PE17:0/17:0 (Avanti Polar Lipids). 1uL of samples were injected into a 1290 Infinity high resolution HPLC coupled with a 6545 Accurate Mass quadrupole time-of-flight (LC-QTOF) (Agilent) via a dual electrospray ionization (ESI) source. Elution of lipids was assessed on a Zorbax Eclipse plus column (Agilent) maintained at 40 °C using an 83 min chromatographic gradient of solvent A (0.2% formic acid and 10 mM ammonium formate in water) and B (0.2% formic acid and 5 mM ammonium formate in methanol/acetonitrile/methyl tert-butyl ether [MTBE], 55:35:10 [v/v/v]). Data acquisition was performed in positive ionisation mode. All samples were processed and analyzed as a single extraction and injection batch. MS quality controls (QCs) were performed by (i) injecting 3 “in-house” QC samples and blanks at the beginning, middle and end of the run and (ii) monitoring the six internal standards spiked in samples for signal intensity, mass-to-charge ratios (m/z) and retention time (RT) accuracies. Mass spectrometry (MS) raw data processing was achieved as previously described using Mass Hunter B.06.00 (Agilent) for peak picking and an in-house bioinformatic script (31) for data processing. The resulting final dataset included 2557 high-quality MS signals, or features, defined by their m/z, RT and signal intensity. Note that several features (adducts) may be annotated for a same unique lipid. Lipid annotation was achieved by MS/MS analysis for 92 significant lipid features that passed the following threshold of significance (using a 2-tailed unpaired Student’s t test with Benjamini-Hochberg correction): a FDR < 0.002 and a fold change (FC) expressed in log2 < 0.5 or > 2.0 for the comparisons (i) G844D vs Ctrl, (ii) G844D vs Ctrl in males and (iii) G844D vs Ctrl in females. In addition, we used data alignment with our in-house database, which contains >500 unique lipids with previously determined MS/MS spectra, for the remaining features. Side chains are indicated when annotated using MS/MS with certitude. When not identified, we only mentioned the total number of carbons and double bounds. The underscore symbol beside acyl chain for several lipids refers to acyl chains for which the sn position remains to be determined.

Unpaired t-test with a Benjamini-Hochberg correction was applied. A threshold of P<0.002 and Fold-change (FC) >2.0 or <0.5 was chosen to select the most discriminant entities between the 2 groups for further validation and annotation using our in-house database APHID (< 5ppm) and the public lipid database LIPID MAPS.

#### Lipoprotein assay

The apolipoprotein B-depleted serum was obtained from the whole serum by precipitating the apolipoprotein B (ApoB) containing lipoproteins with a polyethylene glycol (PEG) solution as described (34). Briefly, whole serum was incubated at 4°C for 20 minutes with a 20% PEG 6000 solution before spinning samples at 10,000 rpm for 30 minutes. The supernatant, which contains the HDL, was collected, and stored at 4°C. Cholesterol and triglycerides from total serum and apoB-depleted serum were quantified using the CHOD-DAOS method and GPO-DAOS method (FujiFilm Wako, Lexington, MA, USA), respectively. Non-HDL cholesterol and triglycerides were obtained by subtraction of the apoB-depleted serum’s results from the total serum’s result.

### Primary mouse hepatocyte culture, radiolabeling, mito-stress assay and treatment

#### Isolation and culture

Mice were anesthetized by isoflurane. Inferior vena cava was cannulated, and liver was perfused by HBSS (GIBCO, 14175-095) with 0.5mM EGTA and digested by low-glucose DMEM (Sigma, D6046) with 1x Penn/Strep (Multicell, 450-201EL), 15uM HEPES (GIBCO, 15630-080) and collagenase IV (Bioshop, COL007) at 100 CDU/mL at flow rate of 1-2mL/min. Dissected liver was placed in digestion medium and released primary hepatocytes filtered through a 70μm Falcon cell strainer (Corning, 352350). Cells were washed in DMEM/F12 (GIBCO, 11320-033) with 1x Penn/Strep and 10% FBS/NCS (Multicell, 098150, 075350) at 50 x g for 2 min at 4°C. Viable hepatocytes were separated in 60% Percoll (Santa Cruz, sc-500790) at 200 x g for 10 min at 4°C. Hepatocytes were washed again. 1.5 million viable hepatocytes were plated as monolayer on collagen-coated 60mm petri dish in isolation medium and attached at 37°C, 5% or 7% CO2 for 2-4 hours. Media was replaced with DMEM (Multicell, 311-425CL) with 1x Penn/Strep for longer incubation and further experiment.

#### Incorporation of [1-^14^C] label into lipids and drug treatment

Use of radioactive compound was approved by the McGill Environmental Health and Safety department and experiments performed at RI-MUHC under protocol (GLEN-0034). [1*-*^14^C]-acetic acid in ethanol (Perkin Elmer, 58 mCi/mmol, 1mCi, NEC084A001MC) was evaporated to dryness under nitrogen and resuspended in a solution of 1M nonradioactive sodium acetate to the indicated specific activity. After the indicated time of attachment, primary hepatocytes were incubated in 2mL of 0.5mM or 1mM [1*-*^14^C]-acetate at 37°C in 5% CO2 incubator with or without 2.5nM insulin (Lonza, BE02-033E20), 5 or 10μM of T0901317 (Cayman Chemical, 70810) for 1-, 3-, 6-, 12- or 24-hours, after which media was collected. Cells were washed and harvested by scraping into PBS (Multicell, 311-425CL) Cell pellet was collected and stored at −20°C for future experiments.

#### BODIPY stain for lipid droplets

Primary hepatocytes seeded on coverslips coated with rat tail collagen type I (Corning, CLS354236) were washed in PBS and fixed by 3% paraformaldehyde (ThermoFisher, 119690010). Cells were then incubated in 5μM BODIPY™ 493/503 (Invitrogen, D3922) for 30 mins and coverslips mounted on slides to enable visualization of lipid droplets.

#### Mito Stress Assay

The Seahorse XF Mito Stress Test Kit (Agilent, 103015-100) was used for this experiment. Primary hepatocytes were seeded in collagen coated XFe96 cell culture microplates (Agilent, 103792-100) at a density of 1.5*10^4^ cells/well in DMEM/F12 containing 10% FBS and 1% Penn/Strep and incubated overnight. Experiment was performed according to Agilent Seahorse XF Mito Stress Test Kit User Guide on the next day, and Mito-Stress assay was run on Seahorse XFe96 Analyzer (Agilent) using Wave software. Medium was removed and cells were lysed and stained by CyQUANT GR dye (Invitrogen, C7026) for viable cell count. Sample fluorescence was measured by fluorescence microplate reader (BMG LabTech). Data was normalized by sample fluorescence value and analysed using Seahorse XF Mito Stress Test Report Generator.

#### Lipid Extraction and column separation

Primary hepatocytes were homogenized using a mini-pestle and extracted by Folch method as previously described. The dried lipid was dissolved in 500μL chloroform and transferred to ISOLUTE^®^ NH2 column (Biotage, 470-0050-B) for separation of each lipid class. Neutral lipids were eluted in 2:1 chloroform:isopropanol; free fatty acids were eluted in 2% acetic acid in diethyl ether; phospholipids were eluted in methanol by gravity (35). Eluents were dried under nitrogen, taken up in Ecolite(+)™ liquid scintillation cocktail (VWR, 88247505), and counted for radioactivity using scintillation counter.

### Statistical analysis

Data analysis was done using the GraphPad Prism software package (version 10.0). Statistical analyses were performed using unpaired, non-parametric t-test, one way or two-way analysis of variance (ANOVA) followed by appropriate correction tests for multiple comparisons. Simple linear regression was performed for correlation analysis. Data are shown as the mean ± standard deviation (SD) and statistical significance was set at P < 0.05. Since we did not observe any significant difference between male and female mice in biochemical analyses, the data from both sexes were combined in the final statistical analysis.

## RESULTS

### Hepatomegaly and histopathological analyses in PEX1-G844D livers

To characterize liver disease progression, liver tissues were collected from mice at age 1 month to 18 months for weight and histology. Progressive relative hepatomegaly was observed in PEX1-G844D homozygotes, depicted by increased liver weight normalized to body weight in mutants across all ages (**Figure 1A)**, whereas liver weight normalized to body weight remained stable in non-mutant littermate controls (7.75 ± 0.837 % at 1-2 months to 19.32 ± 4.12 % at 17-18 months for mutants, *versus* 5.275 ± 0.34 % at 1-2 months to 4.78 ± 0.92 % at 17-18 months for controls).

**Figure 1.**
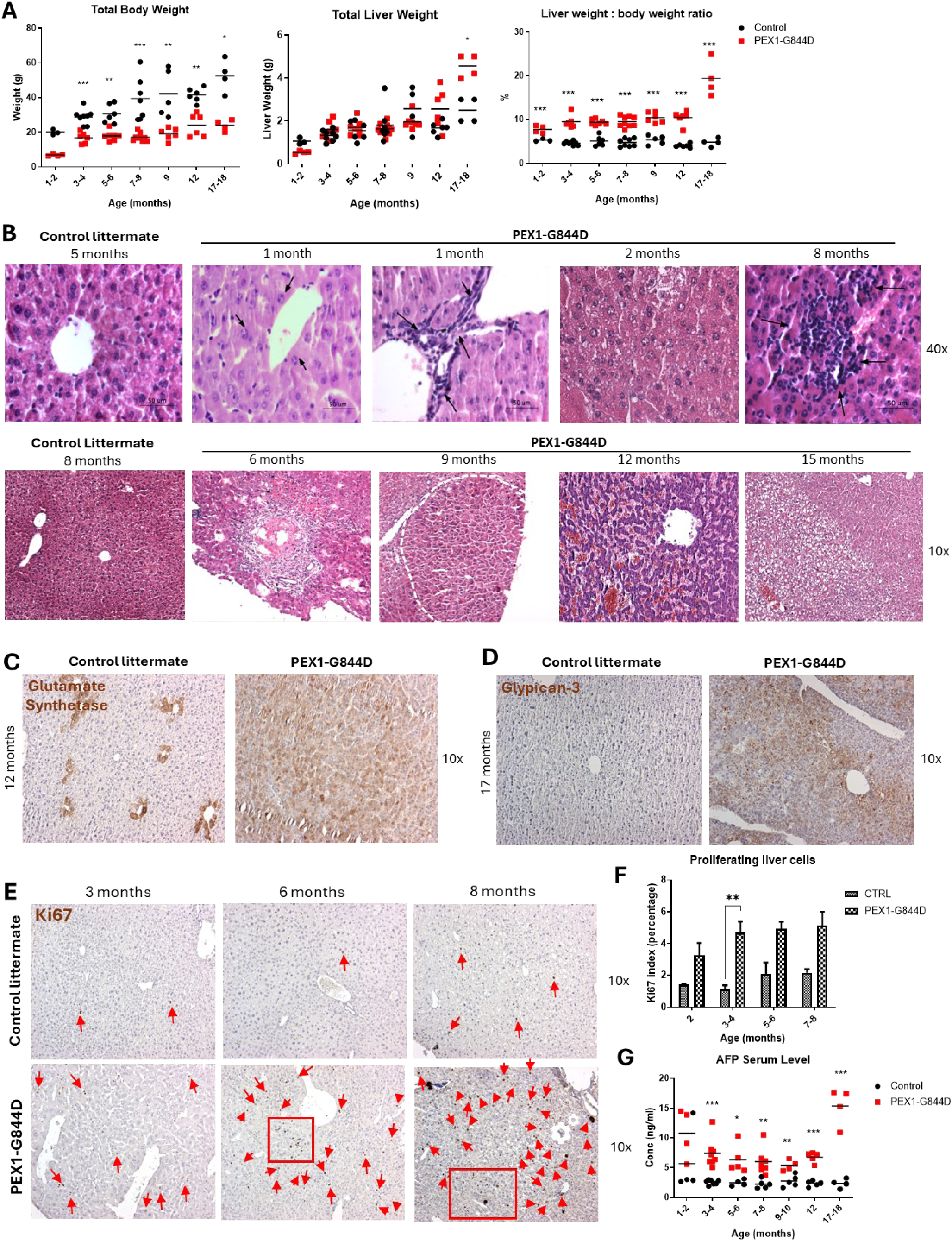
Hepatomegaly and liver disease progression by histology in PEX1-G844D mice. (**A**) Reduced total body weight and increased liver to body weight ratio in PEX1-G844D mutants relative to littermate controls (N=4-6) at all age groups. 1 point represents 1 mouse. (**B**) Hematoxylin and eosin stain (H&E) showed hepatocyte enlargement with prominent nucleoli and ductular reaction (proliferating bile duct with lymphocyte infiltrates, arrows) at 1 month, foamy cell change (2 months), cluster cell-death (6 months), hepatitis (8 months), nodular hyperplasia (dashed line; 9 month), dilated sinusoid with congestion in hepatic tumor (12 months) and distinctive cellular atypia in tumor (15 months) (N=4-6). Top panel: 40x magnification. Bottom panel: 10x magnification. (**C, D**) Immunohistochemistry (IHC) of glutamate synthetase showed diffused positivity in PEX1-G844D mouse liver at age 12 months; glypican-3, a marker of hepatocellular carcinoma (HCC) was present at age 17 months. **(E, F**) IHC for the proliferation marker, Ki67, showed increased number of hepatocytes with Ki67-positive nucleus (red arrows) in PEX1-G844D mutants compared to littermate control at age 2 months, 3-4 months, 5-6 months and 7-8 months (N=3-6). Representative images are shown at age 3, 6 and 8 months. Unpaired student t-test. ** P<0.01; *** P<0.001. (**G**) Serum alpha-fetoprotein (AFP) level was measured across age groups by ELISA. Chronic elevation was observed, with a spike at age 17-18 months. Unpaired student t-test. * P<0.05; ** P<0.01; *** P<0.001.

Liver histology of the lower left lobe visualized by H&E stain is shown in **Figure 1B**. At age 1 month, hypertrophy (hepatocyte enlargement) and ductular reaction (bile duct proliferation) was observed in PEX1-G844D mice. Foamy cells and prominent nucleoli resembling glycogenated nuclei appeared at age 2 months. At age 6 and 8 months, we observed clustered hepatocyte necrosis with lymphocyte infiltration and microvesicular steatosis. At age 9 months, liver nodules started to form, which aggravated significantly to sinusoid dilation that disrupted parenchyma integrity, suggestive of hepatic cancer at age 12 months and 15 months. Diffused positivity for glutamate synthase was observed in liver tumor from PEX1-G844D homozygotes at age 12 months, suggesting the presence of focal nodular hyperplasia or hepatic adenoma (**Figure 1C)**. At age 17 months, liver tumors were positive for Glypican-3, indicating the development of hepatocellular carcinoma (HCC) (**Figure 1D)**.

Higher Ki67 index was observed in livers of PEX1-G844D homozygotes starting at age 2 months (3.11 ± 0.906% for mutants versus 1.409 ± 0.046% for controls), suggesting hepatocyte proliferation (**Figure 1 E, F**). Ki67 index was further increased in PEX1-G844D homozygotes to 4.70 ± 0.675%, 4.916 ± 0.447% and 5.146 ± 0.848% at age 3, 6, and 8 months, respectively. Alpha fetoprotein (AFP), a biomarker indicative of proliferative hepatocytes and hepatic cancer, was consistently elevated in serum of PEX1-G844D homozygotes across all ages and spiked at age 17-18 months, when HCC was evident (**Figure 1G)**. These results suggested continual, progressive hepatocyte hyperplasia in PEX1-G844D homozygotes.

Hepatic glycogen storage was reduced in PEX1-G844D homozygotes at all ages, shown by PAS stain (**Figure 2A)**. Sirius red stain of liver sections (**Figure 2B)** showed a mild increase in collagen deposition around the portal tract, which expanded to periportal parenchyma with time, suggesting subtle development of periportal fibrosis in PEX1-G844D homozygotes. Oil red O stain (**Figure 2C**) for neutral lipids, triglycerides and cholesterol, illustrated increased lipid storage in PEX1-G844D mutants from all age cohorts which was consistent with hepatosteatosis.

**Figure 2.**
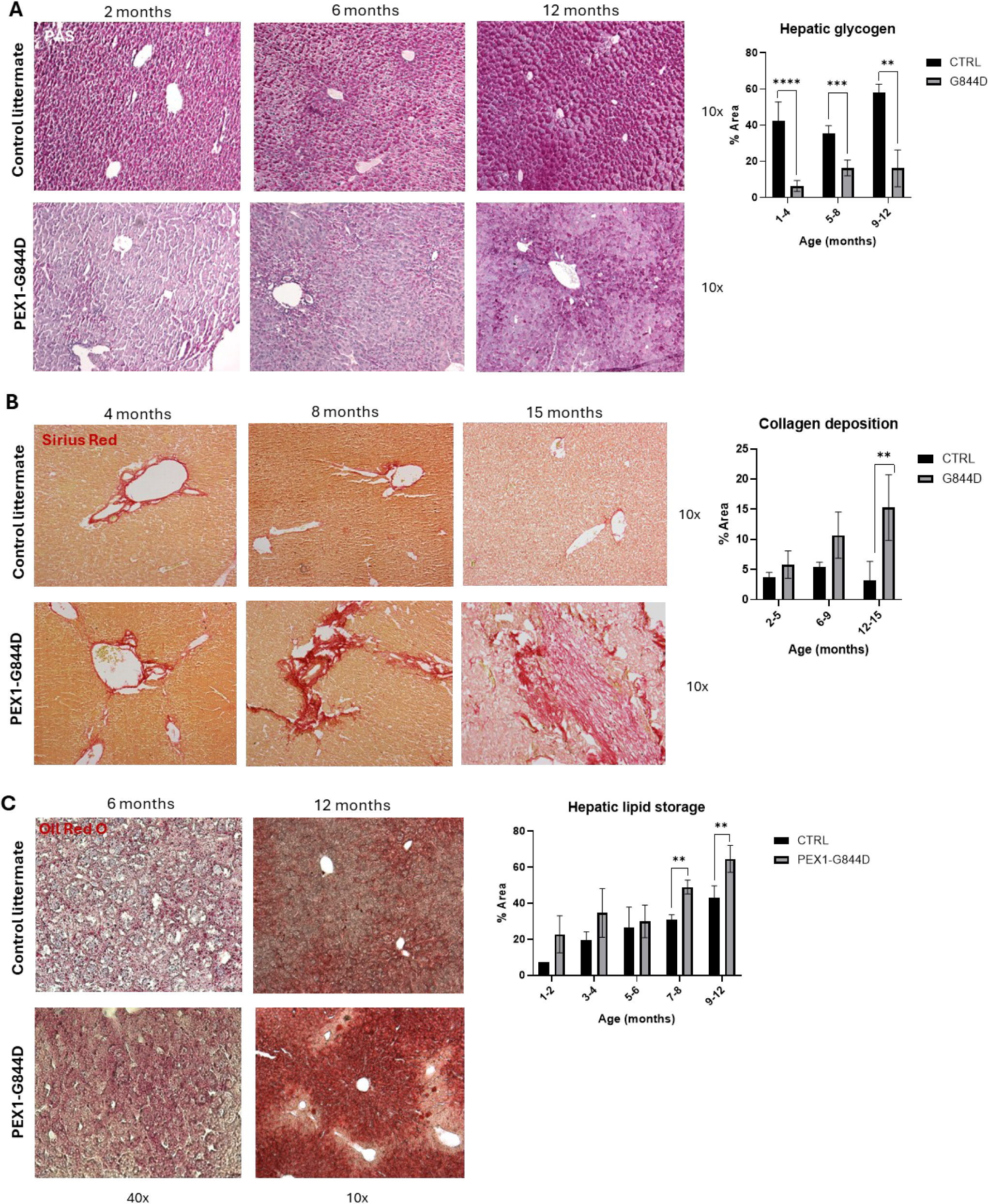
Hepatic glycogen depletion, fibrosis and lipid accumulation in PEX1-G844D mice. (**A**) Periodic-Acid Schiff (PAS) stain for glycogen (purple) and semiquantitative analyses of staining area showed reduced hepatic glycogen storage in PEX1-G844D mice relative to littermate controls across all ages (N=3-5). (**B**) Sirius red stain for collagen (red) semiquantitative analyses showed increasing periportal collagen deposition with time in PEX1-G844D (N=3-5). (**C**) Oil red O stain for neutral lipids (triglycerides and cholesterol, red) and semiquantitative analyses for staining area indicated accumulation of lipid droplets in liver of PEX1-G844D mutants (N=3-5) across all ages suggestive of hepatosteatosis. Top panel: 40x magnification; bottom panel: 10x magnification. Representative images for PAS, Sirius red and oil red O are shown. Semiquantitative analyses of staining area showed significant reduction in hepatic glycogen storage and accumulation of hepatic lipid storage and collagen deposition in G844D mutants relative to controls. Unpaired student t-test. ** P<0.01; *** P<0.001.

### Defective peroxisome assembly, abnormal hepatocyte ultrastructure, and secondary mitochondrial defect in PEX1-G844D livers

Immunofluorescent microscopy was used to compare peroxisome number and peroxisomal import functions in primary hepatocytes from PEX1-G844D homozygotes and littermate controls. Peroxisome membrane protein 70 (PMP70) was used as a peroxisome marker to visualize peroxisome numbers and size. There were few to no peroxisomes in PEX1-G844D liver sections compared to littermate controls (**Figure 3A**). Catalase, a peroxisomal matrix protein, was used to visualize peroxisomal import. When import is intact, catalase primarily localizes inside peroxisomes and co-localizes with PMP70. In PEX1-G844D hepatocytes there was diffuse cytosolic catalase distribution, indicating defective import into any residual peroxisomes.

**Figure 3.**
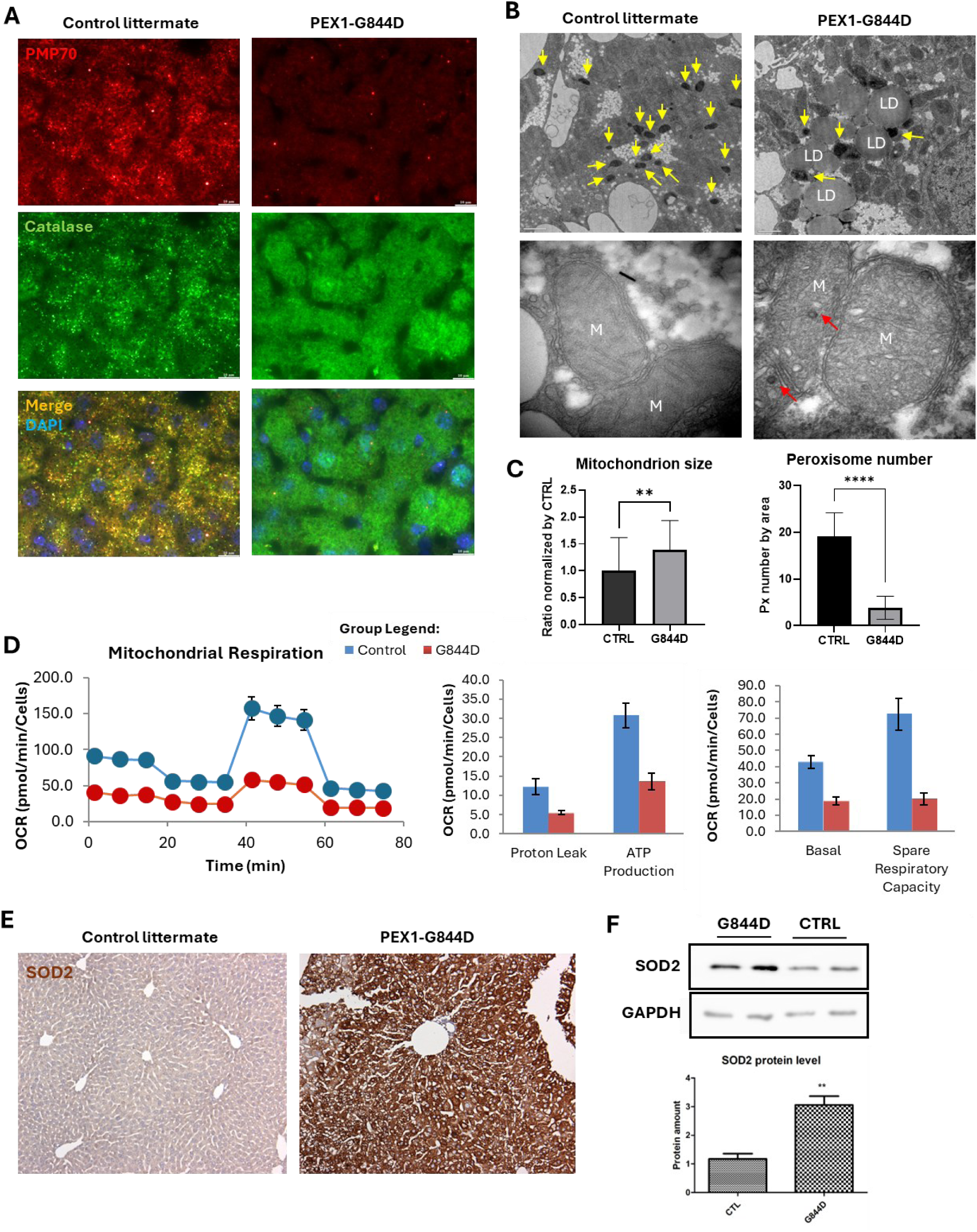
Defective peroxisome assembly, abnormal hepatocyte ultrastructure and secondary mitochondrial defect, in PEX1-G844D livers. (**A**) Liver immunofluorescence on paraffin-embedded tissue revealed few PMP70 positive peroxisomes and cytosolic catalase localization, suggesting defective peroxisome import in PEX1-G844D mice (N=3 per genotype). Representative images shown at age 2 months. (**B**) Transmission electron microscopy (TEM) of 3,3′-diaminobenzidine stained PEX1-G844D mouse hepatocytes from 3-month-old mice showed fewer electron-dense peroxisomes (yellow arrows) compared to littermate controls with accumulation of lipid droplets (LD) (top panels, 2,900x magnification) and enlarged mitochondria (M) with dilated cristae and electron dense particles (red arrows) (bottom panel, 11,000 x magnification). Findings were consistent in N=4 mice per genotype. (**C**) Average mitochondrial surface area and peroxisome number normalized by area. Unpaired student t-test. * P<0.05; ** P<0.01. (**D**) PEX1-G844D mutants showed enlarged mitochondrial and reduced peroxisome relative to controls. Unpaired student t-test. ** P<0.01; **** P<0.0001. (**D**) Seahorse Mito Stress assay revealed reduced basal mitochondrial respiration and ATP production in primary hepatocytes from PEX1-G844D homozygotes relative to littermate controls at age 3-4 months. Findings were consistent in N=3 per genotype. (**E**) Liver immunohistochemistry revealed increased superoxide dismutase 2 (SOD2) in PEX1-G844D mice (N=3-5 per genotype). Representative images are shown at age 2 months. (**F**) Higher SOD2 protein level was detected in PEX1-G844D liver relative to controls by immunoblot analysis (N=4 per genotype at age 2-3 months). Unpaired student t-test. * P<0.05.

To observe any structural abnormalities in subcellular organelles consequent to peroxisome dysfunction, hepatocyte ultrastructure was visualized by transmission electron microscopy. Catalase was stained by 3,3′-Diaminobenzidine to detect import-competent peroxisomes. In PEX1-G844D homozygotes, there was a significant reduction in the number of peroxisomes compared to littermate controls (**Figure 3B**). These residual functional peroxisomes were enlarged, and the DAB stain was irregular. There was accumulation of lipid droplets in PEX1-G844D mutant tissues. We also observed enlarged mitochondria (**Figure 3B, C)** with dilated mitochondrial cristae and abnormal electron dense particles in the matrix, which is a typical feature reported in patients with mitochondrial disorders (36). Enlarged mitochondria were also observed in retinal tissue in this model (22).

To investigate real-time mitochondrial respiratory function *in vitro,* Seahorse Mito Stress Assay was performed on primary hepatocytes harvested from PEX1-G844D homozygotes and littermate controls at age 2-3 months. We observed an approximately 50-70% reduction in maximal respiration, proton leak and ATP production in PEX1-G844D mutants compared to controls (**Figure 3D)**. There was also an approximate 60% reduction in basal oxygen consumption and spare respiratory capacity in mutants. Together, this evidence suggested a defective mitochondrial electron transport chain and reduced ATP production in the PEX1-G844D liver. Given that mitochondria are also required to metabolise reactive oxygen species generated from mitochondrial respiration, we investigated mitochondrial oxidative stress by visualizing superoxide dismutase 2 (SOD2), a mitochondrial anti-oxidative enzyme, using IHC. We observed an overall increase in SOD2 across liver sections at age 2-4 months (**Figure 3E**). Immunoblotting confirmed higher SOD2 protein levels in mutant liver lysate (**Figure 3F**). Together, these results indicated early mitochondrial dysfunction and increased mitochondrial oxidative stress consequent to the primary peroxisome deficiency.

### PEX1-G844D protein levels, peroxisome metabolites, and abnormal liver functions

Although the PEX1-G843D allele is known to be degraded (37), leading to decrease in its partner proteins PEX6 and PEX5, the murine PEX1-G844D protein was shown to be stable and at normal level in retinal tissue (22) and liver of our mouse model using immunoblot (**Supplemental Figure 1A**). In addition, its binding partner PEX6, and their ligand PEX5 (the peroxisome enzyme receptor) protein in mutant livers were also comparable to controls, suggesting that the murine missense protein is not degraded.

Lipid analyses were performed on flash-frozen liver tissue from PEX1-G844D mutant mice and littermate controls from weaning (3 weeks) to 37 weeks of age (**Supplemental Figure 1B**). At age 10-16 weeks, very-long chain fatty acids, C26:0 and C24:0, and branched-chain fatty acids, pristanic and phytanic acids were elevated in mutants, which were 2.5-fold, 2.2-fold, 58-fold and 51-fold of average control values, respectively. Hepatic levels of C27 bile acid precursors, peaked at age 10-16 weeks, averaging 282- and 550-fold for DHCA and THCA, respectively, and remained elevated in PEX1-G844D mutants across the ages. In contrast, the C24 mature bile acid, cholic acid, was reduced to 9-41% of average control values in mutant liver. Hepatic levels of PE 22:6 plasmalogens and total plasmalogens were also reduced to 17% and 50% of average control values in mutant liver, respectively (data not shown).

Serum liver function tests were performed in mice from 3 to 37 weeks of age (**Supplemental Figure 2**). There was significant elevation of ALP levels in mutants across ages by 2.5-3.2-fold of average control values. ALT and AST were elevated in mutants by 6.8-fold and 2.2-fold of control levels at 24-30 weeks of age, respectively. There was a mild reduction in albumin in mutants to 87% of control levels at age 31-37 weeks. Triglycerides were reduced to 42-58% of control at age 10-16 weeks. Asymptomatic hypoglycemia was observed in PEX1-G844D mice across ages, showing 51-62% of average control values. There was no significant difference in cholesterol levels between mutants and littermate controls (3.893 ± 0.892 mmol/L versus 3.08 ± 1.42 mmol/L, see Supplemental Figure 2).

### Abnormal lipidomic profiles in blood and liver of PEX1-G844D mice

To investigate the hepatic lipid profile in PEX1-G844D, untargeted lipidomic analysis was performed on serum and flash-frozen liver tissue (**Figure 4, Supplemental Tables 2** and **3**). In PEX1-G844D serum, besides the 7.52-fold elevation of C26:0 lyso-phosphatidylcholine (PC C26:0_0:0) relative to littermate controls, there was 2-5-fold elevation of other very-long-chain PC, long-chain PC, and 3.78-fold increase of 18:3 cholesteryl ester. Consistent with the serum biochemical profile, triglycerides were reduced to 2-29% of average control values in PEX1-G844D mutants. In addition, they showed deficiency in sphingomyelins (SM d44:3, d18:1/22:1, d17:1/16:0) and ether-PC plasmalogens (PCO-), which were 18-44%, and 25-38% of control, respectively (**Figure 4A, B, Supplemental Table 2**). Deficiency of these membrane lipid species recapitulates the systemic dyslipidemia reported in ZSD patients (38).

**Figure 4.**
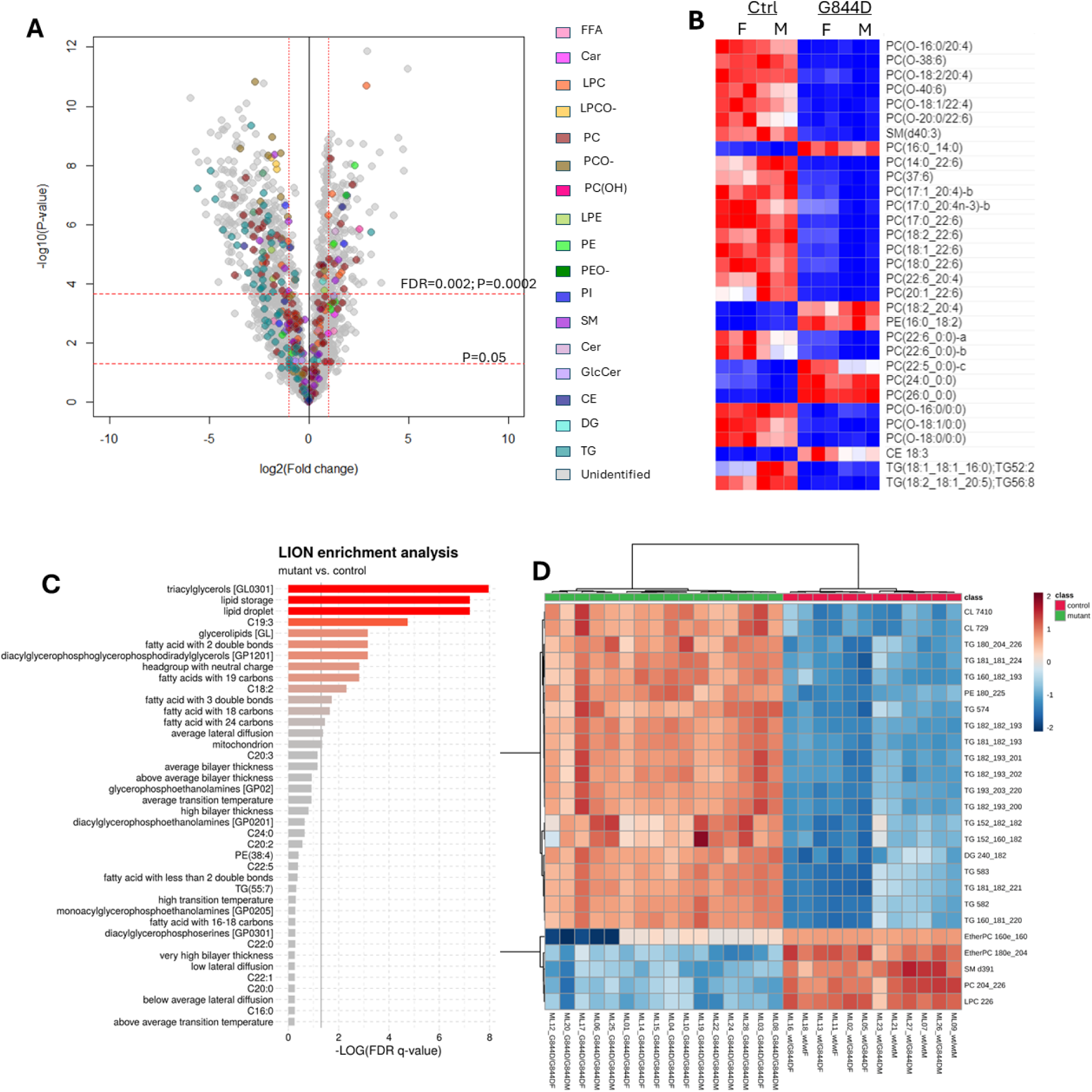
Serum fatty acid deficiency and altered hepatic lipid profile in PEX1-G844D mice. (**A**) Volcano plot of MS/MS validated lipid classes showed reduction of various lipid species among triglycerides (TG), cholesteryl esters, sphingomyelins, phosphatidylcholines (PC) and ether phosphatidylcholines in serum of PEX1-G844D mutants relative to controls at age 2.5 months (N=8 per genotype). (**B**) Heat map of fold-change (FC) of serum lipid species annotated by MS/MS with FC>2.0 or FC<0.5 at P value =<0.002 revealed major systemic deficiency of PC, ether-PC (PCO) and TG in PEX1-G844D mutants. This included deficiency of DHA (PC 22:6) containing PC. Accumulation of circulating VLCFA-PC (24:0, 26:0) was observed in PEX1-G844D mutants at age 2.5 months as expected (N=6 per genotype). (**C**) Lipid Ontology Term (LION) analysis demonstrated hepatic accumulation of triacylglycerols and increased lipid storage in PEX1-G844D mice at age 6 months. (**D**) Heat map of top 25 hepatic lipid species with most significant differential amount between mutants (N=16) and control (N=12) revealed hepatic buildup of triglyceride (TG)s, cardiolipins, phosphatidylethanolamines, with deficiency of ether PCs, sphingomyelins and unsaturated PCs in PEX1-G844D mouse model.

In contrast to serum, PEX1-G844D liver tissue presented a different lipid profile. Membrane lipid species, such as ether-PC (16:0_16:0, 18:0_20:4) and ether-PE plasmalogens, sphingomyelin (d39:1) and PC (18:3_22:6, 20:4_22:6), were also deficient in PEX1-G844D homozygotes (20%, 80%, 40% and 9-37% of control, respectively). However, there were prominent accumulations of triglycerides and cardiolipin compared to littermate controls (**Figure 4C)**. Lipid ontology term (LION) enrichment analysis suggested a build-up of lipid species stored as lipid droplets in PEX1-G844D mouse liver, consistent with the increased lipid droplets and micro-steatosis observed by histology (**Figure 4D)**. In contrast, long-chain fatty acids and monounsaturated fatty acids were decreased (**Supplemental Table 3)**, which may compromise the functions of intracellular organelles and cell membranes.

### Investigation of hypoglycemia and disrupted hepatic lipid homeostasis

We hypothesized that hypoglycemia in PEX1-G844D homozygotes would lead to lower circulating levels of insulin by physiological response, and hypoinsulinemia would sequentially disrupt downstream lipid homeostasis in liver. To investigate this, we measured circulating insulin by immunoassay, which showed approximately 59-63% reduction in PEX1-G844D serum compared to littermate controls in both fasting (5-6 hours) and fed (*ad libitum*) conditions (**Figure 5A**).

**Figure 5.**
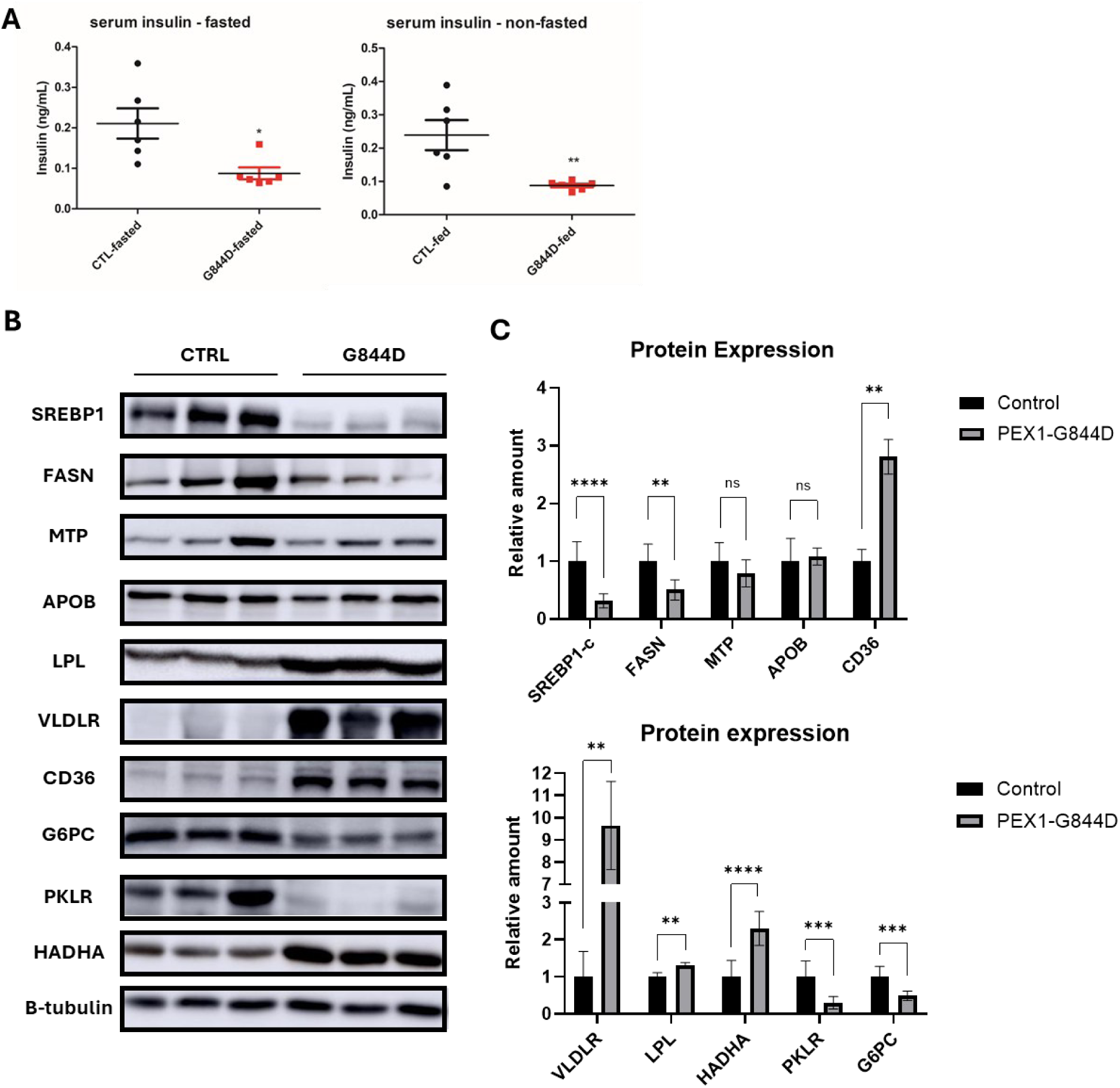
Hypoinsulinemia and dysregulated hepatic lipid homeostasis in PEX1-G844D mice. (**A**) ELISA quantification of serum insulin showed hypoglycemia in PEX1-G844D mutant mice relative to littermate controls at age 2.5 months in both fasted and fed *ad-libitum* state (N=6 per genotype). (**B, C**) Immunoblots quantified by densitometry revealed lower levels of proteins involved in hepatic *de novo* lipogenesis (SREBP1, FASN), glycolysis (PKLR) and glycogenolysis (G6PC), and increased amounts of proteins involved in hepatic lipid/lipoprotein uptake (VLDLR and CD36), lipoprotein breakdown (LPL) and fatty acid oxidation (HADHA) in liver of PEX-G844D mutants compared to littermate controls at age 2.5 months (N=6 per genotype). Unpaired student t-test. * P<0.05; ** P<0.01; *** P<0.001; ns: not significant.

To investigate downstream metabolic pathways in liver, RT-qPCR was performed to determine the transcript levels of key regulatory genes involved in lipid and carbohydrate homeostasis (**Table 1**). Expression of genes involved in hepatic uptake of circulating lipids, was prominently increased. Transcripts of *CD36*, *Vldlr* and *Lpl*, genes regulating hepatic uptake and clearance of free FA and VLDL, were elevated by 18.23-fold, 24.22-fold and 6.18-fold, respectively, relative to littermate controls. In addition, the expression level of *Hadha*, involved in hepatic FA oxidation, was also increased by 2.21-fold. In contrast, average gene expression levels of *Srebf-1* and *Fasn*, key regulators of *de novo* hepatic FA synthesis, were reduced by 3.42-fold and 1.54-fold, respectively, in PEX1-G844D homozygotes compared to controls. Genes involved in synthesizing triacylglycerol for hepatic storage (*Gpam)* and packing triacylglycerol into very-low-density lipoprotein (VLDL) for extrahepatic secretion (*Apob* and *Mttp*) were insignificantly increased PEX1-G844D livers. These results suggested that increased hepatic uptake of circulating fatty acids was the major contributor to observed hepatosteatosis in PEX1-G844D mice.

**Table 1.**
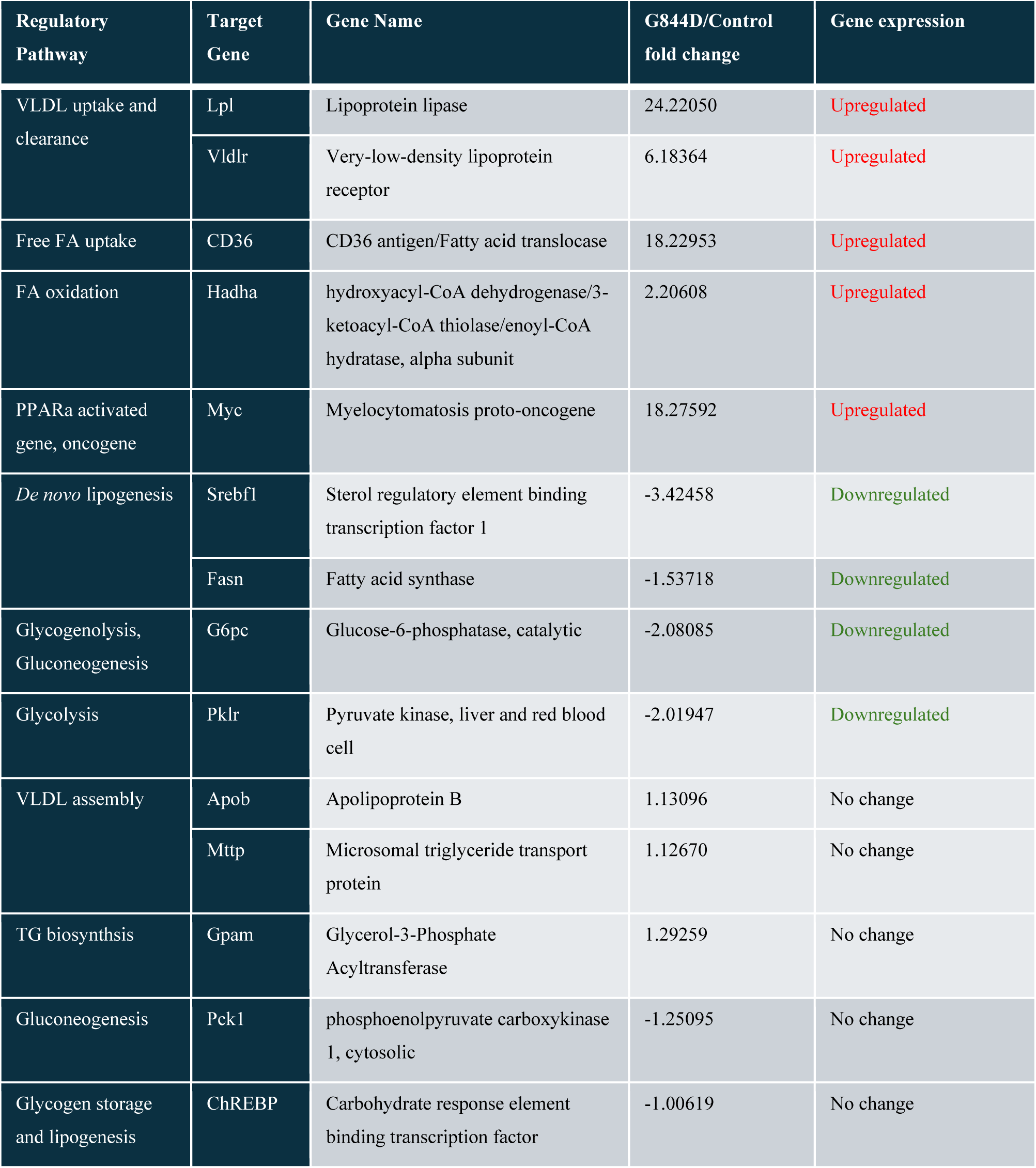
Quantitative expression of genes involved in hepatic lipid and glycogen metabolism. RT-qPCR analysis of selected key regulatory genes involved in disrupted metabolic pathways in PEX1-G844D livers. N=6 per genotype. Threshold of fold change > (+/-) 1.5 are considered as a change in gene expression.

Although G844D homozygotes were hypoglycemic, the expression level of *Chrebp*, which promotes hepatic glycogen synthesis and lipogenesis upon activation was not significantly reduced (−1.006-fold) (**Table 1**). In addition, expression of genes participating in glycolysis and glycogenolysis (*G6pc* and *Pklr*) was downregulated by 2.08-fold and 2.02-fold compared to littermate controls, whereas expression of a gene involved in gluconeogenesis (*Pck1*) was unchanged. Taken together, these observations could explain the consistent hypoglycemia and subsequent chronic hypoinsulinemia in PEX1-G844D mice.

Finally, an 18.27-fold increase of *Myc*, together with other downstream targets of PPARα (*Cd36* and *Lpl*), confirmed activation of the PPARα signalling pathway in PEX1-G844D homozygotes. PPARα pathways are normally activated in fasting states to drive catabolism of lipids and glucose for energy production (39, 40). Natural activators of PPARα include unsaturated fatty acids, eicosanoids, as well as pristanic and phytanic acids (41).

To confirm the gene expression studies, we evaluated the corresponding protein levels by immunoblot on liver tissue homogenates (**Figure 5B**). Consistent to transcript levels, we observed reduced amounts of SREBP1, FASN, G6P and PKLR proteins measuring 32%, 51%, 49% and 30% of control, respectively (**Figure 5C**). We also confirmed increased amounts of CD36, VLDLR, LPL and HADHA proteins, measuring 2.8-fold, 9.6-fold, 1.3-fold and 2.3-fold of control, respectively. Protein levels of MTP and APOB were unaltered (0.79-fold and 1.08-fold of control), suggesting normal hepatic assembly of VLDL for extrahepatic trafficking.

To assess the consequence of enhanced hepatic FA uptake, serum lipoproteins were precipitated and separated into high-density lipoproteins (HDL) and non-HDL. Triglycerides and cholesterols in each fraction were measured (**Figure 6A)**. We observed an overall 61-64% reduction in total and non-HDL associated triglycerides. This observation supported increased VLDL uptake from circulation into liver. The non-significant reduction of HDL-associated triglycerides might also contribute to hypotriglyceridemia in PEX1-G844D mutants. Although total and HDL-associated cholesterol was normal, we observed a 3.5-fold elevation of non-HDL associated cholesterol in PEX1-G844D mutants. This observation implies an abnormal cholesterol trafficking in PEX1-G844D mice.

**Figure 6.**
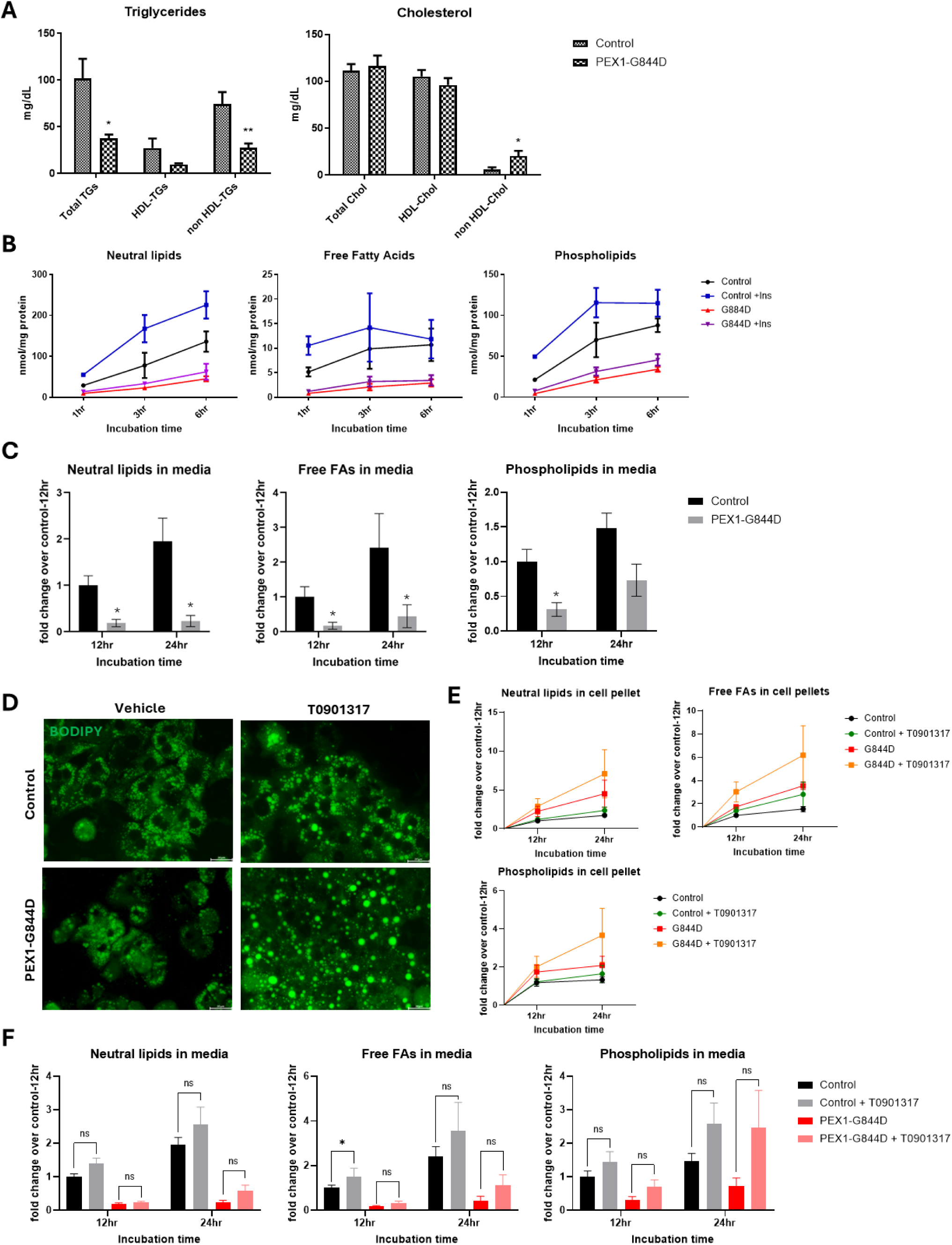
Disrupted lipoprotein-trafficking and reduced *de novo* fatty acid synthesis in PEX1-G844D mice. **(A)** Lipoprotein precipitation analysis showed reduced lipoprotein-associated triglycerides, in particular, non-HDL triglycerides and increased non-HDL cholesterol in PEX1-G844D mouse serum relative to control at age 2.5 months (N=6 per genotype). (**B**) Incubation of ^14^C radiolabelled acetate with primary hepatocytes showed lower hepatic *de novo* fatty acid synthetic rate in all lipid classes (neutral lipids, free fatty acids and phospholipid) in PEX1-G844D homozygous hepatocytes relative to controls at age 2-3 months (N=3). Insulin supplementation increased the rate of fatty acid biosynthesis in primary hepatocytes of both genotypes. (**C**) Reduced ^14^C labelled lipids in media of PEX1-G844D mutant hepatocytes after 12- and 24- hours incubation with ^14^C-acetate to recapitulate *in vivo* systemic lipid deficiency observed in the PEX1-G844D mouse model. Unpaired student t-test. * P<0.05; ** P<0.01. (**D**) BODIPY fluorescence stain for lipid droplets showed increased size of lipid droplets in both control and PEX1-G844D mutant hepatocytes after 48-hour incubation with T0901317 (N=3 per genotype). (**E**) Incorporation of ^14^C-radiolabel into each lipid class suggested greater retention and accumulation of newly synthesized lipid contents after 12 and 24 hours of T0901317 incubation in PEX1-G844D mutant hepatocytes compared to those of controls (N=3 per genotype). (**F**) Measurement of ^14^C-radiolabelled lipids in cell media confirmed the effect of T0901317 to promote hepatic fatty acid secretion in both PEX1-G844D mutant and control primary hepatocytes (N=3 per genotype). Unpaired t-test. * P<0.05; ns: not significant.

### Assessing hepatic *de novo* lipogenesis *in vitro*

To assess hepatic *de novo* fatty acid biosynthesis, primary hepatocytes harvested from PEX1-G844D homozygotes and non-mutant littermate controls at age 8-10 weeks were incubated in 0.5mM [1-^14^C] radiolabeled acetate for 1, 3 and 6 hours (**Figure 6B)**. Relative amounts of [1-^14^C] radiolabel incorporated into each lipid class was measured. We observed lower levels of neutral lipids, free FAs and phospholipids synthesized over 1-, 3-, and 6-hour incubation times in PEX1-G844D homozygotes, suggesting a reduced rate of hepatic *de novo* lipogenesis (DNL) relative to littermate controls. To determine if added insulin recovers hepatic *de novo* lipogenesis in PEX1-G844D mice, insulin supplementation at 2.5ng/mL was applied to hepatocytes. Although insulin stimulated hepatic DNL by increasing neutral lipids and phospholipids after 3- and 6-hour incubation in both mutant and control hepatocytes, it did not recover hepatic DNL in G844D homozygotes to normal levels. These findings support the stimulatory effect of insulin on hepatic *de novo* lipogenesis in this model.

To assess the rate of fatty acid secretion *in vitro*, primary hepatocytes were incubated with 1mM of [1-^14^C] radiolabeled acetate for 12 and 24 hours. Cell media was collected for lipid extraction and detection of ^14^C-radiolabel lipids secreted into media. There was a 69-89% reduction in the amount of radiolabeled neutral lipids, free FAs, and phospholipids secreted from PEX1-G844D mouse hepatocytes compared to non-mutant controls (**Figure 6C)**. These observations are consistent with reduced serum lipids determined by our previous experiments.

### Liver X Receptor Agonist enhanced DNL and lipid secretion

T0901317 is a synthetic liver X receptor (LXR) agonist developed to treat atherosclerosis. Despite its athero-protective effect (42, 43), T0901317 was not approved by the FDA due to its lipogenic side-effect (44–46). In liver, LXR is a transcription factor upregulated by insulin receptor activation (i.e. high insulin), leading to downstream activation of genes involved in hepatic lipid synthesis (e.g. *SREBF1*, *FASN*, *ACC*) (47). In animal models, activation of LXR by T0901317 upregulates hepatic FA synthesis genes to increase plasma levels of triglycerides and phospholipids (48–51). Therefore, we hypothesized that treatment with T0901317 in PEX1-G844D mice will improve systemic FA deficiency by enhancing DNL and extrahepatic lipid trafficking.

We treated primary hepatocytes harvested from PEX1-G844D homozygotes and littermate controls with 10uM T0901317 or vehicle (0.1% DMSO) for 12- and 24-hour with 1mM [1-^14^C] radiolabeled acetate. Lipid extracted from cell pellets and cell media was analyzed for radiolabel-incorporation into each lipid class (**Figure 6E, F**). We observed a 3.2-fold and 3.4-fold increase in neutral lipid and free FA secretion in PEX1-G844D hepatocytes after 24-hour treatment with T0901317 compared to vehicle-treated PEX1-G844D hepatocytes. Secretion of phospholipids was recovered in PEX1-G844D relative to control cells after 24-hour T0901317 treatment (2.471 ± 2.219-fold in mutants *versus* 2.585 ± 1.241-fold in controls). In control hepatocytes, the amount of lipid secretion was also enhanced by T0901317 relative to vehicle treatment (2.585 ± 1.241-fold *versus* 1.479 ± 0.693-fold). However, the accumulation of all lipid classes in PEX1-G844D hepatocytes was exaggerated by T0901317 after 12- and 24-hours incubation (2.6-fold to 4.1-fold for neutral lipids, 2.3-fold to 4-fold for free FAs, 1.56-fold to 2.75-fold for phospholipids). This observation was corroborated by BODIPY fluorescence stain for neutral lipids in hepatocytes treated with T0901317 *versus* DMSO for 48 hours, which showed an increased number and size of lipid droplets in T0901317-treated PEX1-G844D and control hepatocytes compared to vehicle-treated hepatocytes (**Figure 6D**). Together, these findings suggest that although T0901317 induced hepatic DNL and lipid secretion in PEX1-G844D homozygotes, it also exaggerated hepato-steatosis, which will aggravate hepatic disease progression in PEX1 deficiency.

## DISCUSSION

Liver disease is one of the major phenotypes in the PEX1-G844D mouse model of mild ZSD. This may be related to dramatic reduction of peroxisomes in liver, whereas near normal numbers were present in retina and fibroblasts from G844D mutant mice (22). These observations suggest loss of sub functional peroxisomes in liver which may underly the early onset of hepatic dysfunction and hepatomegaly as well as liver disease progression in PEX1-G844D mouse model and ZSD patients. Detailed characterization of hepatopathy progression in this model provided several mechanisms for the major findings of hepatic cancer, hepatosteatosis, systemic lipid deficiency and glycogen depletion diagramed on **Figure 7**.

**Figure 7.**
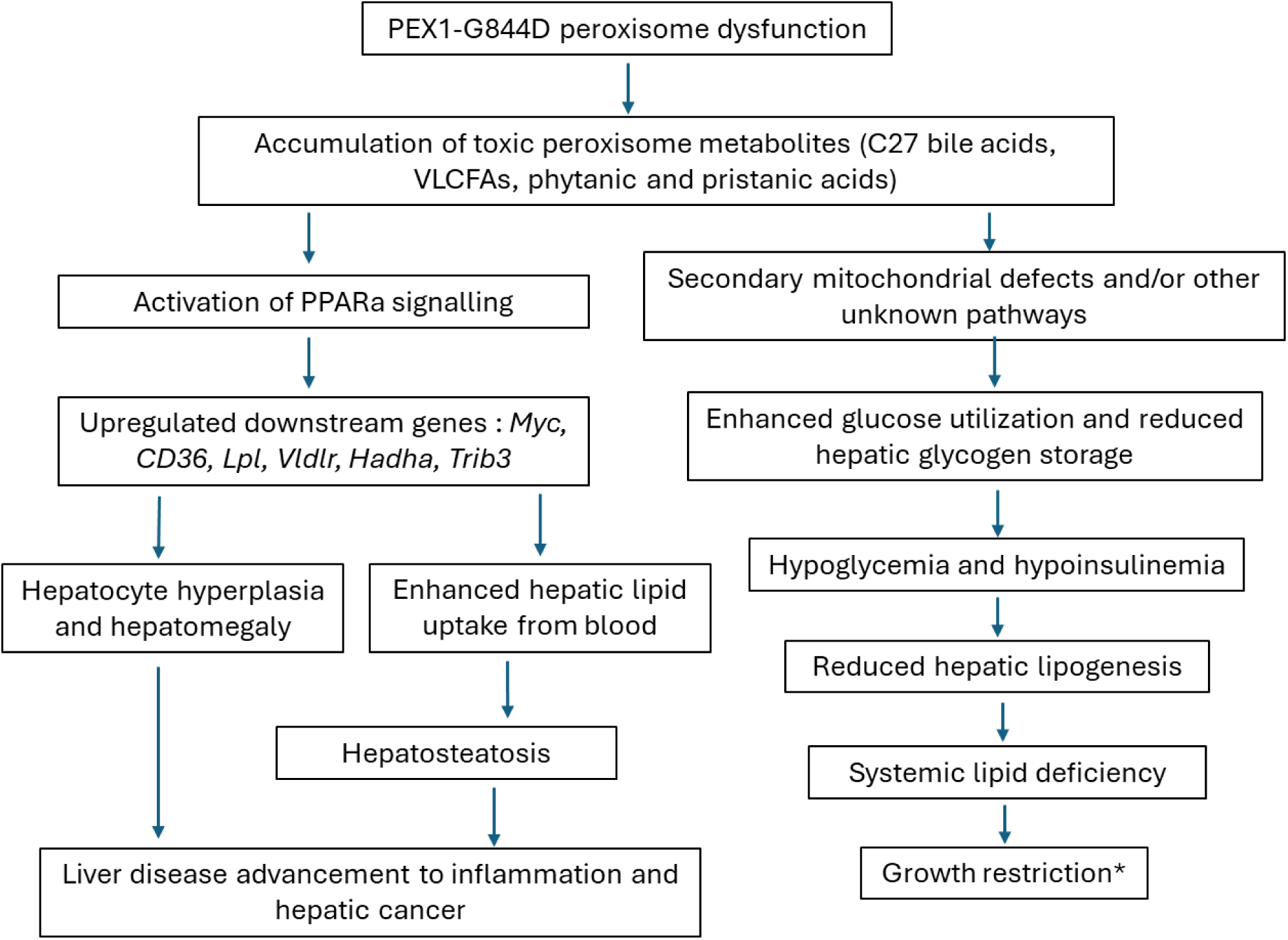
Proposed pathway disturbances underlying liver disease progression in PEX1-G844D mice. *Deficiency of mature bile acid may also contribute to growth restriction.

### Tumor development in PEX1-G844D mice

Previous literature reported early hepatopathy in PEX1-G844D mice, including hepatomegaly, elevated liver enzymes, steatosis, hyperplasia, cholestasis and fibrosis (18, 19). Preliminary characterization of our model showed similar findings (17). In this manuscript, we showed that liver histology in PEX1-G844D mutants is constantly abnormal. Structural anomalies are apparent in the first month of age, and gradually progress to hepatic tumor and suspected hepatocellular carcinoma at age 15-18 months. Chronic elevation of AFP levels and progressively increasing Ki67 index suggest ongoing hepatocyte hyperplasia, which likely contributes to hepatomegaly and the development of hepatic tumors in this model.

The ongoing hepatic proliferation may be correlated with increased expression of the *Myc* oncogene. We suggest that *Myc* is upregulated by PPARα activation. As reported in other PEX1-G844D murine models (18, 19), increased PPARα signalling is likely due to an accumulation of branched chain fatty acid ligands, phytanic and pristanic acids, also found in our model (**Supplemental Figure 1B**). Our observations also complement previous literature showing PPARα activation and hepatic tumor formation in other PBD mouse models with *Pex2, Alb-Pex5, Acox1* and *Dbp* knockout, whereas *Acox1/Pparα* double knockout mice did not develop liver cancer (52, 53). Together, these findings support the role of PPARα activation in promoting hepatocarcinogenesis in a non-cirrhotic background in peroxisome-deficient mouse models. However, in human ZSD, hepatocellular carcinoma usually arises from a cirrhotic background (16, 52, 54) implying a different etiology.

### Mitochondrial defects

We found increased mitochondrial oxidative stress and defective mitochondrial respiratory function in our PEX1-G844D hepatocytes, which is consistent with findings in liver of *Alb-Pex5^- /-^* mice and a PEX1-G844D mouse model congenic on C57BL/6 background (19, 55). In rat hepatoma cells, it was shown that accumulation of C27 bile acid metabolites (DHCA and THCA) uncouples oxidative phosphorylation, inhibits respiratory chain complexes, and enhances production of mitochondrial reactive oxygen species (ROS) (56). In another study of mitochondrial respiratory function in liver tissue from liver specific *Alb-Pex5*^-/-^ and *Mfp2*^−/−^ (defect in multifunctional protein, a β-oxidation enzyme) mice, C27 bile acid intermediates and other peroxisome metabolites were not solely responsible for hepatic mitochondrial anomalies (57). Supplementation of our PEX1-G844D mice with cholic acid, which normalizes C27 bile acid intermediates, reduced mitochondrial oxidative stress demonstrated by reduced SOD2 levels (unpublished results). We therefore speculate that mitochondrial oxidative stress and respiratory defects in PEX1-G844D hepatocytes could be due to the accumulation of toxic C27 bile acid metabolites.

### Glycogen metabolism

Our results indicate chronic reduction of hepatic glycogen storage over lifetime. With gene expression analysis, we observed downregulation of genes involved in glycolysis, gluconeogenesis, glycogenolysis and glycogen synthesis (*Pklr, Pck1*, *G6pc* and *Gsy2,* respectively). Reduction of hepatic glycogen was reported in the PEX1-G844D mice congenic on NMRI background from age 1-6 months (18) and *Alb-Pex5*^-/-^ mice(58). In both reported models, gene expression showed downregulation of gluconeogenesis with increased glycolysis or enhanced glycolytic enzyme activity (58). This latter observation implied potential post-transcriptional or post-translational mechanisms of glycolytic induction. Co-regulation of peroxisomal and carbohydrate metabolism was demonstrated in *pex2* and *pex6* mutant drosophila, leading to enhanced glycolysis, reduced glycogen and susceptibility to glucose deprivation by hepatic activation of AMP-activated protein kinase (AMPK) and suppression of PGC-1α pathways (59). However, significant alteration of AMPK and PGC-1α pathways were absent in our PEX1-G844D mice (data not shown), implicating alternate pathways coordinating peroxisome biogenesis and carbohydrate metabolism. It was previously suggested that secondary mitochondrial respiratory chain energy defect resulted in reliance on glucose metabolism for energy and drive glucose utilization(19). Overall, these findings suggested that impaired glycogen synthesis and/or increased glycolysis consequent to peroxisomal defects could be the major contributor to reduced glycogen storage in peroxisome deficient mice.

### Lipid homeostasis

Hepatic and systemic lipidomic profiles were not previously investigated in any PEX1-G844D mouse model. Our PEX1-G844D mice showed an overall reduction in circulating triglycerides, phosphatidylcholine (PC), and sphingomyelins (SM). This finding corroborates with the dyslipidemia in ZSD patients (38), which also showed a systemic lipid deficiency of PC, phosphatidylethanolamine (PE), and sphingolipids, all membrane lipids. Considering that these are main constituents of cellular membranes, trafficking vesicles and signalling molecules (60), systemic fatty acid deficiency in the PEX1-G844D mouse model and patients may contribute to the multi-systemic involvements of brain, liver and growth restriction. In comparison, liver tissue of PEX1-G844D mice showed a different lipid profile. We found that accumulation of triglycerides and cholesterols in G844D, with increased hepatic uptake of circulating fatty acids and lipoproteins, underlies systemic lipid deficiency.

Hypo-insulinemia consequent to hypoglycemia observed in PEX1-G844D homozygotes contributes to downregulation of genes involved in hepatic lipogenesis confirmed by reduced SREBP1 and FASN protein levels (61, 62). In addition, we showed increased fatty acid synthesis in G844D hepatocytes after insulin supplementation, suggesting that the reduction of hepatic lipogenesis in G844D homozygotes is a physiological response to chronic hypo-insulinemia. Overall, we propose a cycle of hypoglycemia-hypoinsulinemia-downregulated hepatic *de novo* lipogenesis-systemic lipid deficiency which further drives glucose combustion and energy depletion in our ZSD mouse model and possibly in patients. On the other hand, accumulation of peroxisome metabolites results in liver-activated PPARα signalling, which in turn upregulates downstream pathways including hepatic lipids uptake, hepatic VLDL clearance, and FA oxidation(63, 64). Together, these contribute to increased hepatic lipid storage, hepatosteatosis and systemic fatty acid deficiency in PEX1-G844D mice. Though LXR agonist, T0901317, improves systemic FA deficiency in PEX1-G844D mice via promoting hepatic *de novo* lipogenesis and enhancing hepatic FA secretion, exacerbation of hepatosteatosis becomes a concern. The combination of LXR agonist and pharmacological inhibition of hepatic lipid uptake may prevent hepatosteatosis and provide desired treatment outcome.

In summary, our PEX1-G844D mouse model exhibited an early onset hepatic pathology that progressed to focal necrosis, hepatosteatosis, increased collagen deposition and ultimate hepatic cancer. Our characterization complements the progression seen in equivalent human disease, in which a subset of patients showed hepatomegaly, cholestasis, fibrosis, cirrhosis and hepatocellular carcinoma (52). Our in-depth hepatic phenotyping in the PEX1-G844D mouse model provides robust therapeutic endpoints for preclinical trials. These endpoints include preserving normal hepatic lobular architecture and glycogen amount, and preventing steatosis and tumor formation. Deteriorating histopathology over time reveals an early therapeutic window at age 4-6 weeks, which showed minimal abnormalities of hepatocyte enlargement, bile duct proliferation and depletion of hepatic glycogen storage. The pathways identified underlying disrupted lipid homeostasis and carbohydrate metabolism should provide potential therapeutic targets for candidate drug therapies to prevent hepatopathy in ZSD.

## Author contributions

LC designed the experiments, collected and processed tissues, performed all mouse-related experiments, carried out data analyses and interpretations and wrote the manuscript under supervision by NB. HC assisted with experimental design. CA established and maintained our PEX1-G844D mouse colony, collected data, prepared and visualized samples to provide initial preliminary data with MH. WC established the genotyping protocol for PEX1-G844D mouse colony and assisted with mouse liver perfusion. EDP performed LC-MS/MS lipid analyses. ZHG and VHN assisted with histology and ultrastructure interpretation. CD, MR and EN performed LC-MS/MS untargeted lipidomic analysis. DC and DR performed lipoprotein analysis. JGH performed microarray analysis. NB conceptualized the study, assisted with manuscript writing and reviewing.

## Financial support

This research is funded by a Canadian Institutes of Health Research Project Grant to NB and institutional grant RI-MUHC New Direction Research Award to NB. LC was awarded salary support from an RI-MUHC studentship award.

## Acknowledgement

We would like to thank Lyne Joyal for assistance with interpretation of hepatocyte ultrastructure, Johanne Ouelette, Jeannie Mui, S. Kelly Sears and staffs from the Facility of Electron Microscopy Research (FEMR) at McGill University for assistance with sample preparation and transmission electron microscopy usage. We would like to thank Dr. Frederic M Vaz (Laboratory Genetic Metabolic Diseases, University of Amsterdam, Netherland) for his measurements on peroxisome metabolites in liver tissues.

## Conflict of interest

There is no conflict of interest

## List of Abbreviations

AFP: Alpha-fetoprotein
ALT: Alanine Aminotransferase
ALP: Alkaline Phosphatase
APOB: Apolipoprotein B
AST: Aspartate Aminotransferase
ATP: adenosine triphosphate
BA: Bile acids
cDNA: complimentary Deoxyribonucleic Acid
ChREBP: Carbohydrate Response Element Binding Protein
COX IV: cytochrome oxidase subunit IV
CTRL: control
CYP7A1: cholesterol 7α-hydroxylase
DAB: 3,3′-Diaminobenzidine
DHCA: 3α,7α,12α-dihydroxycholestanoic acid
DNL: De novo lipogenesis
ELISA: enzyme-linked immunosorbent assay
FA: Fatty acids
FASN: Fatty acid synthase
FC: Fold change
FDR: False Discovery Rate
FXR: Farnesoid X receptor
GC/MS: Gas Chromatography/Mass Spectrometry
G6PC: Glucose-6-Phosphatase, catalytic
HADHA: hydroxyacyl-CoA dehydrogenase trifunctional multienzyme complex subunit alpha
HCC: Hepatocellular carcinoma
HDL: High-density lipoprotein
LC-MS/MS: Liquid chromatography-tandem mass spectrometry
LDL: Low-density lipoprotein
LFT: Liver Function Tests
LPL: Lipoprotein lipase
LXR: Liver X Receptor
LPC/lyso-PC: lysophosphatidylcholine
MTP: Microsomal transfer protein
OCR: Oxygen Consumption Rate
PBD: peroxisome biogenesis disorders
PC: phosphatidylcholine
PCO: ether-phosphatidylcholines
PE: phosphatidylethanolamine
PEO: ether-phosphatidylethanolamine
PEX: Peroxin
PKLR: Pyruvate Kinase Liver and Red blood cell
PL: plasmalogen
PMP: peroxisome membrane proteins
PPAR: peroxisome proliferator-activated receptor
PTS: peroxisome targeting signal
PUFA: Polyunsaturated fatty acids
qRT-PCR: quantitative real-time polymerase chain reaction
SD: Standard Deviation
SKL: Serine-Lysine-Leucine
SM: Sphingomyelins
SOD2: Superoxide dismutase 2
SREBP1: Sterol Regulatory Element Binding Protein 1
TEM: Transmission Electron Microscopy
TG: Triglycerides
THCA: 3α,7α,12α-trihydroxycholestanoic acid
VLCFA: Very-long-chain fatty acids
VLDL: Very-low-density lipoproteins
ZSD: Zellweger Spectrum Disorder

## Notes

### Competing Interest Statement

The authors have declared no competing interest.

### Summary of Updates

This version of the manuscript has been revised to update author information.

