## Supplemental Table and Supplemental Figure for "Longitudinal study of liver disease progression in the PEX1-Gly844Asp mouse model of mild Zellweger Spectrum Disorder"

**Supplemental data**

**Supplemental Table 1 List of primers used for quantitative real-time PCR**

| Genes | Forward (5' to 3') | Reverse (5' to 3') |
|---|---|---|
| *B2m* | CTGGTCTTTCTGGTGCTTGTC | GCAGTTCAGTATGTTCGGCTT |
| *Gusb* | CAGGGTTTCGAGGCAGCAATG | ACCCAGCCAATAAAGTCCCG |
| *Srebf1-c* | GGAAGCTGTCGGGGTAGCG | GCATAGGGGGCGTCAAACAG |
| *Fasn* | CCAAGCAGGCACACACAATG | CAGTGTTCGTTCCTCGGAGT |
| *Mttp* | GTCACACAACTGGCCTCTCA | CCTCTCTGTTGACCCGCATT |
| *Apob* | TCCAAAGAGGCCAGTCAAGC | GGACTGCCTGTTCTCAACCA |
| *Cd36* | GGCTGTGTTTGGAGGCATTC | CCACGTCATCTGGGTTTTGC |
| *Vldlr* | ACCTGTTCCTGTCCCAATGG | TCACTGTAAGTCACAGGAGTTGAAGTAC |
| *Lpl* | CGAGAGCGAGAACATTCCCT | CCTCTCGATGACGAAGCTGG |
| *Gpam* | CAACACCATCCCCGACATC | GTGACCTTCGATTATGCGATCA |
| *Hadha* | CAGGCAAGGGAAGGTCATCA | AAGGCGTCCAGCTTCTTAGG |
| *Pklr* | CAGCATCATTGCCACCATCG | GACTCCAGTGCGTATCTCGG |
| *G6pc* | TGGGCATCAATCTCCTCTGG | AATACGGGCGTTGTCCAAAC |
| *Pck1* | TGCAGAACACAAGGGCAAGAT | TTTGCCGAAGTTGTAGCCGA |
| *Chrebp* | AAGTTGCTATGCCGGGACAA | AGGTTTCCGGTGCTCATCTG |
| *Myc* | TACGGAGTCGTAGTCGAGGT | GGATTTCCTTGGGCGTTGG |
| *Slc27a5* | GCGGTACTTGTGTAACGTCC | ATTTGCCCGAAGTCCATTGC |

**Supplemental Table 2A. 19 lipid species increased in serum of PEX1-G844D mice**

| Annotation | Adduct | Compound | FDR | FC (abs) | Regulation |
|---|---|---|---|---|---|
| CE 18:3 | Fragment | POS:368.3447@50.05 | 0.000002 | 3.78 | up |
| CE 18:3 | [M+NH4]+ | POS:663.5965@50.04 | 0.000004 | 3.20 | up |
| Cer(d18:1/24:0) | [M+Na]+ | POS:671.6199@38.79 | 0.00002 | 2.53 | up |
| GlcCer(d18:1/24:0) | [M+H]+ | POS:811.6901@36.56 | 0.00100 | 2.37 | up |
| PC(16:0_14:0) | [M+H]+ | POS:705.5316@21.69 | 0.0000003 | 2.13 | up |
| PC(16:0_18:3) | [M+H]+ | POS:755.5480@21.49 | 0.00009 | 5.32 | up |
| PC(18:2_0:0)-a | [M+H]+ | POS:519.3338@8.01 | 0.00001 | 2.01 | up |
| PC(18:2_20:4) | [M+H]+ | POS:869.6886@37.24 | 0.000001 | 4.58 | up |
| PC(20:2_18:2) | [M+H]+ | POS:809.5949@24.81 | 0.00001 | 3.05 | up |
| PC(22:5_0:0)-c | [M+H]+ | POS:569.3486@9.37 | 0.00025 | 2.88 | up |
| PC(24:0_0:0) | [M+H]+ | POS:607.4585@17.25 | 0.000002 | 2.26 | up |
| PC(26:0_0:0) [M+H]+ | [M+H]+ | POS:635.4896@20.80 | 0.00000001 | 7.52 | up |
| PC(36:4(OH)) | [M+Na]+ | POS:797.5560@19.82 | 0.00001 | 5.91 | up |
| PE(16:0_18:1) | [M+H]+ | POS:717.5308@26.92 | 0.00003 | 2.42 | up |
| PE(16:0_18:2) | [M+H]+ | POS:715.5161@24.06 | 0.0000004 | 4.99 | up |
| PE(18:0_18:2) | [M+H]+ | POS:743.5480@28.04 | 0.000002 | 3.71 | up |
| SM(d18:1/24:0)-c | [M+H]+ | POS:814.6946@36.24 | 0.00036 | 2.55 | up |
| SM(d18:2/24:0) | [M+H]+ | POS:812.6780@34.29 | 0.00009 | 2.82 | up |
| TG(18:1_18:1_22:4);TG58:6 | [M+NH4]+ | POS:951.8258@63.14 | 0.00002 | 8.70 | up |

**Supplemental Table 2B. 51 lipid species reduced in serum of PEX1-G844D mice**

| Annotation | Adduct | Compound | FDR | FC (abs) | Regulation | Annotation | Adduct | Compound | FDR | FC (abs) | Regulation |
|---|---|---|---|---|---|---|---|---|---|---|---|
| CE 22:6 | [Dimer+Na]+ | POS:1415.1506@48.85 | 0.00007 | 0.22 | down | PI(16:0_20:4) | [M+C2H5NH3]+ | POS:903.5821@18.92 | 0.000003 | 0.44 | down |
| CE 22:6 | Fragment | POS:160.1251@48.86 | 0.00004 | 0.11 | down | SM(d17:1/16:0) | [M+H]+ | POS:688.5525@18.97 | 0.00017 | 0.44 | down |
| PC(14:0_22:6) | [M+H]+ | POS:777.5314@18.83 | 0.000001 | 0.11 | down | SM(d18:1/22:1) | [M+H]+ | POS:784.6461@29.12 | 0.00003 | 0.18 | down |
| PC(16:0_22:6) | [M+H]+ | POS:805.5651@22.22 | 0.00024 | 0.19 | down | SM(d40:3) | [M+H]+ | POS:782.6301@25.96 | 0.0000003 | 0.31 | down |
| PC(17:0_20:4n-3)-b | [M+H]+ | POS:795.5785@24.90 | 0.00027 | 0.33 | down | TG(16:0_16:0_22:6);TG54:6 | [M+NH4]+ | POS:895.7634@51.97 | 0.000003 | 0.21 | down |
| PC(17:0_22:6) | [M+H]+ | POS:819.5780@24.11 | 0.00001 | 0.10 | down | TG(16:0_18:2_20:5);TG54:7 | [M+C2H5NH3]+ | POS:921.7776@46.59 | 0.00003 | 0.28 | down |
| PC(17:1_20:4)-b | [M+H]+ | POS:793.5627@21.80 | 0.00001 | 0.17 | down | TG(16:0_18:2_20:5);TG54:7 | [M+NH4]+ | POS:893.7483@46.64 | 0.00001 | 0.11 | down |
| PC(18:0_22:6) | [M+H]+ | POS:833.5953@26.03 | 0.00005 | 0.15 | down | TG(18:1_16:0_20:4);TG54:5 | [M+NH4]+ | POS:897.7806@55.20 | 0.00001 | 0.24 | down |
| PC(18:1_22:6) | [M+H]+ | POS:831.5791@22.81 | 0.00001 | 0.14 | down | TG(18:1_16:0_20:4);TG54:5 | [M+C3H7NH3]+ | POS:939.8262@55.23 | 0.00077 | 0.29 | down |
| PC(18:2_22:6) | [M+H]+ | POS:829.5627@20.32 | 0.00002 | 0.26 | down | TG(18:1_16:0_22:6);TG56:7 | [M+C3H7NH3]+ | POS:963.8256@52.33 | 0.000001 | 0.02 | down |
| PC(20:1_22:6) | [M+H]+ | POS:859.6097@26.48 | 0.000005 | 0.18 | down | TG(18:1_16:0_22:6);TG56:7 | [M+NH4]+ | POS:921.7799@52.35 | 0.000001 | 0.03 | down |
| PC(20:4_20:3) | [M+H]+ | POS:831.5778@21.96 | 0.00001 | 0.20 | down | TG(18:1_16:0_22:6);TG56:7 | [M+CH3NH3]+ | POS:935.7948@52.35 | 0.000002 | 0.04 | down |
| PC(22:6_0:0)-a | [M+H]+ | POS:567.3333@7.98 | 0.00008 | 0.23 | down | TG(18:1_17:0_18:2);TG53:3 | [M+NH4]+ | POS:887.7952@62.01 | 0.00063 | 0.20 | down |
| PC(22:6_0:0)-b | [M+H]+ | POS:567.3338@8.32 | 0.00012 | 0.20 | down | TG(18:1_18:1_16:0);TG52:2 | [M+NH4]+ | POS:875.7957@65.57 | 0.00047 | 0.15 | down |
| PC(22:6_20:4) | [M+H]+ | POS:853.5629@19.74 | 0.000002 | 0.08 | down | TG(18:1_18:1_22:4);TG58:6 | [M+C2H5NH3]+ | POS:979.8571@61.71 | 0.00008 | 0.20 | down |
| PC(37:6) | [M+H]+ | POS:791.5475@20.45 | 0.00002 | 0.15 | down | TG(18:2_16:0_20:4);TG54:6 | [M+C3H7NH3]+ | POS:937.8094@50.06 | 0.00001 | 0.25 | down |
| PC(O-16:0/0:0) | [M+H]+ | POS:481.3545@9.97 | 0.0000003 | 0.27 | down | TG(18:2_18:1_20:5);TG56:8 | [M+NH4]+ | POS:919.7635@46.94 | 0.0000001 | 0.13 | down |
| PC(O-16:0/20:4) | [M+H]+ | POS:767.5836@25.17 | 0.0000001 | 0.28 | down | TG52:5 | [M+NH4]+ | POS:869.7486@49.37 | 0.00063 | 0.19 | down |
| PC(O-18:0/0:0) | [M+H]+ | POS:509.3853@11.69 | 0.0000004 | 0.32 | down | TG52:6 | [M+C2H5NH3]+ | POS:895.7632@44.49 | 0.00115 | 0.24 | down |
| PC(O-18:1/0:0) | [M+H]+ | POS:507.3696@10.39 | 0.000001 | 0.33 | down | TG53:2 | [M+C2H5NH3]+ | POS:917.8417@71.28 | 0.00023 | 0.12 | down |
| PC(O-18:1/20:4) | [M+H]+ | POS:793.5991@25.71 | 0.000002 | 0.38 | down | TG54:7 | [M+NH4]+ | POS:893.7481@47.38 | 0.00001 | 0.05 | down |
| PC(O-18:1/22:4) | [M+H]+ | POS:821.6290@28.61 | 0.0000003 | 0.25 | down | TG54:8 | [M+NH4]+ | POS:891.7334@44.26 | 0.00002 | 0.07 | down |
| PC(O-18:2/20:4) | [M+H]+ | POS:791.5858@22.80 | 0.0000003 | 0.38 | down | TG56:7 | [M+NH4]+ | POS:921.7795@50.05 | 0.00002 | 0.27 | down |
| PC(O-20:0/22:6) | [M+H]+ | POS:847.6451@32.48 | 0.000003 | 0.19 | down | TG56:8 | [M+C2H5NH3]+ | POS:947.7949@46.93 | 0.000002 | 0.21 | down |
| PC(O-38:6) | [M+H]+ | POS:791.5832@24.33 | 0.00000001 | 0.15 | down | TG58:6 | [M+NH4]+ | POS:951.8260@55.70 | 0.00017 | 0.21 | down |
| PC(O-40:6) | [M+H]+ | POS:819.6145@28.31 | 0.0000002 | 0.09 | down | | | | | | |

**Supplemental Table 2 Legend. Dyslipidemia was identified by untargeted lipidomic analysis on serum samples of PEX1-G844D mice.** In PEX1-G844D mutants, 70 annotated lipids analyzed by MS/MS were either increased by 2-fold **(A)** or reduced to 50% **(B)** of levels in control littermates (false discovery rate (FDR) $< 0.002$, N=6 per genotype). Unpaired t-test with Benjamini-Hochberg correction. FC, fold change.

**Supplemental Table 3. Hepatic lipidomic profile of PEX1-G844D mice.**

| Class | mutant | control | mutant/control |
|---|---|---|---|
| Cer | 9627 | 3429 | 2.8 |
| EtherOxPC | 30 | 34 | 0.9 |
| EtherPC | 7 | 35 | 0.2 |
| EtherPE | 219 | 279 | 0.8 |
| FA | 29 | 27 | 1.1 |
| FAHFA | 1 | 1 | 1.0 |
| GM3 | 1 | 0 | 2.4 |
| HexCer | 328 | 232 | 1.4 |
| LPE | 139 | 73 | 1.9 |
| MGDG | 75 | 17 | 4.4 |
| PA | 143 | 293 | 0.5 |
| PC | 19312 | 16614 | 1.2 |
| PE | 9980 | 5579 | 1.8 |
| PEtOH | 24 | 27 | 0.9 |
| PG | 89 | 78 | 1.1 |
| PI | 20 | 13 | 1.5 |
| PMeOH | 11 | 24 | 0.5 |
| PS | 314 | 198 | 1.6 |
| ACar | 21 | 14 | 1.5 |
| BMP | 6 | 5 | 1.2 |
| CE | 1478 | 1833 | 0.8 |
| cholesterol | 3 | 2 | 1.6 |
| CL | 4206 | 1060 | 4.0 |
| DG | 1620 | 2146 | 0.8 |
| LPC | 486 | 548 | 0.9 |
| SM | 535 | 1243 | 0.4 |
| TG | 11420 | 6236 | 1.8 |

**Supplemental Table 3 Legend.** Altered hepatic lipid profile by lipid class in PEX1-G844D mice compared to non-mutant controls. Elevation and reduction in the amount of each lipid class is presented by fold change in mutant relative to controls.

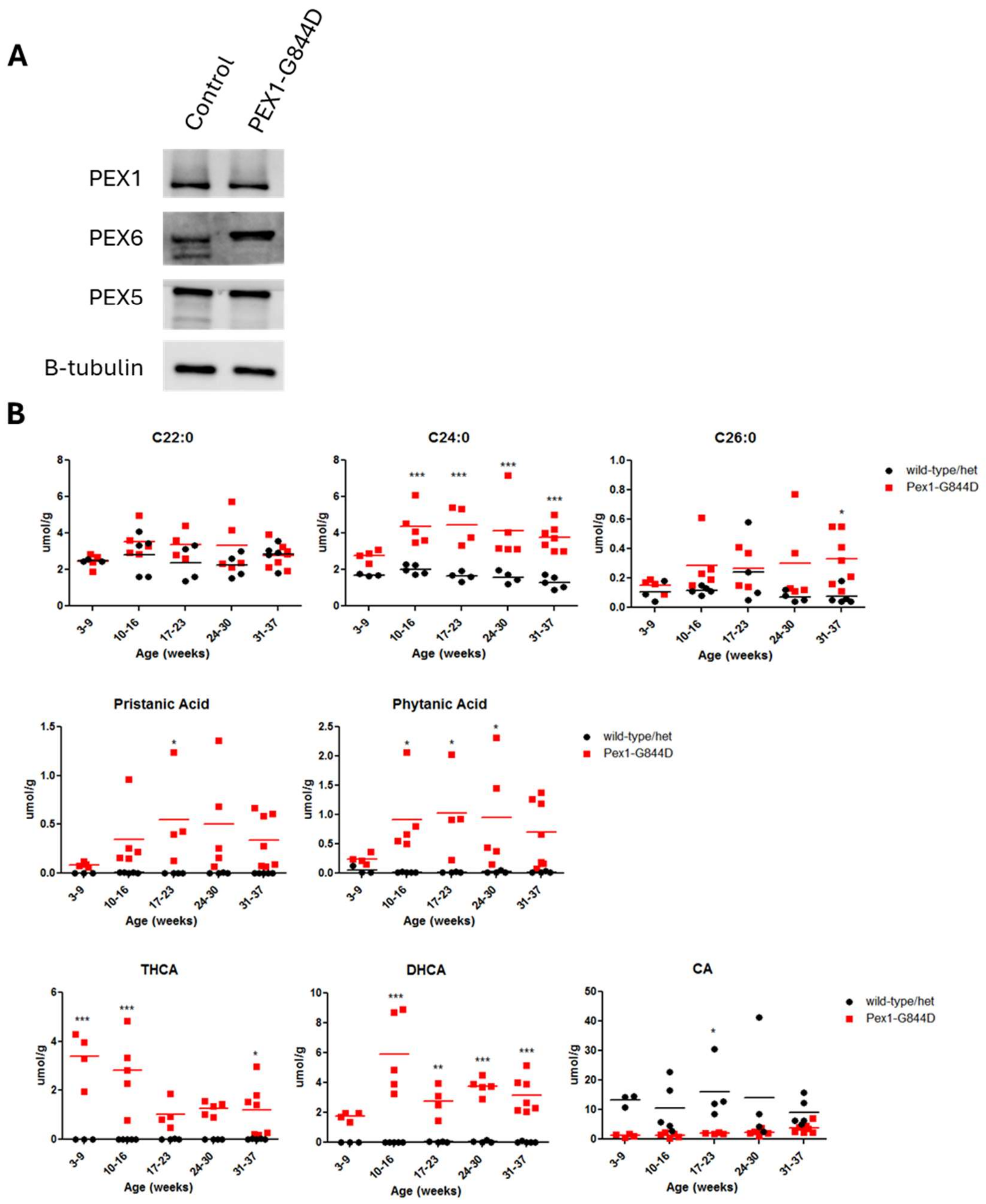


**Supplemental Figure 1. Peroxisomal proteins and metabolites in livers.** (**A**) Immunoblotting of liver lysates showed equal PEX1 protein levels between PEX1-G844D and littermate controls. PEX6 and PEX5 levels were also normal (N=3). Representative images from 2-month-old mice are shown. (**B**) LC-MS/MS analysis on flash-frozen liver tissue showed elevated C24:0 and C26:0 VLCFAs, pristanic and phytanic acids, and C27 bile acid precursors (DHCA and THCA) in PEX1-G844D homozygotes relative to littermate controls (N=3-7). Reduction of mature cholic acid (CA) was observed at all ages. Samples were assessed at age 3-9 weeks, 10-16 weeks, 17-23 weeks, 24-30 weeks and 31-37 weeks. * P<0.05; ** P<0.01; *** P<0.001 (unpaired t-test).

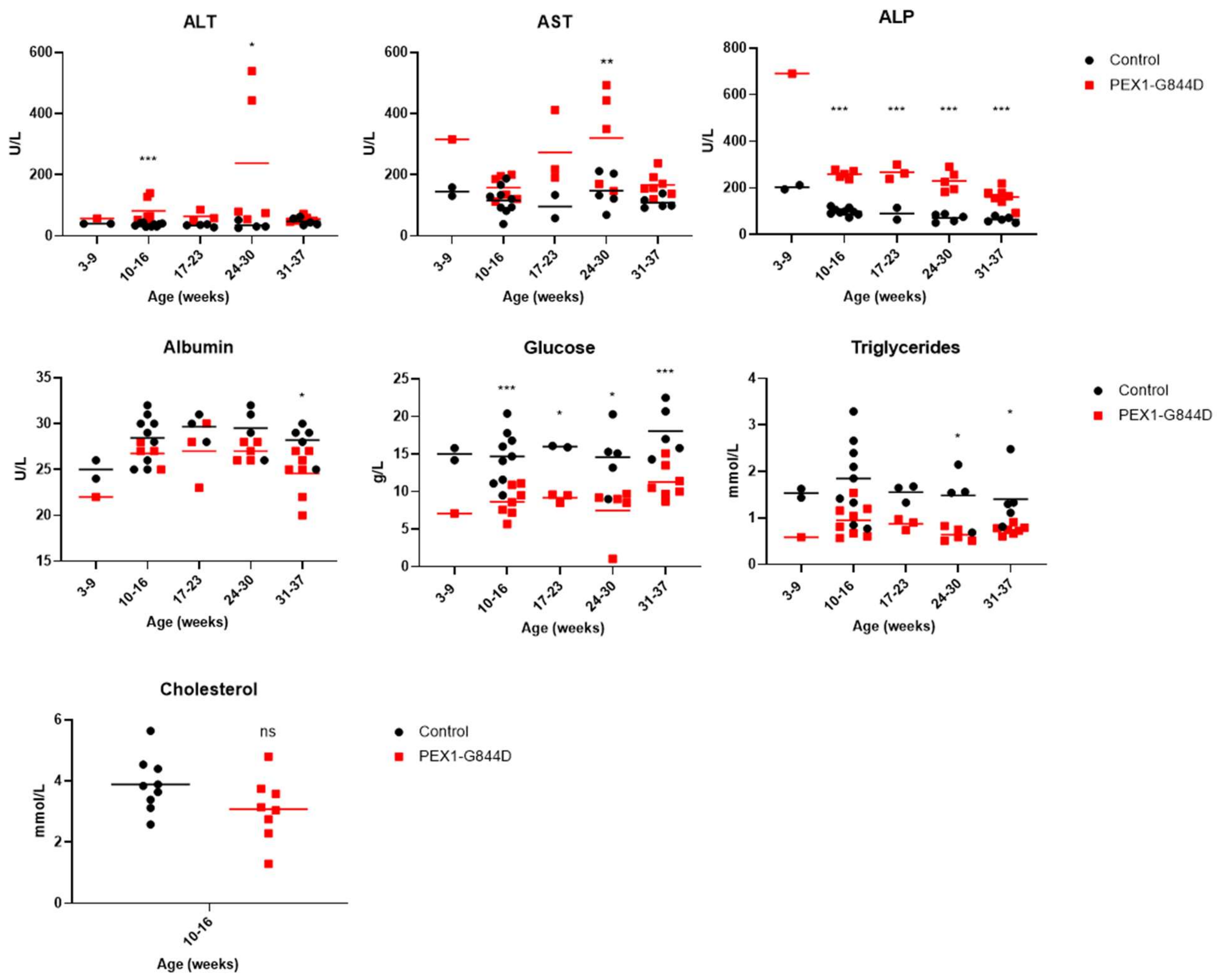


**Supplemental Figure 2. Liver function tests in PEX1-G844D mice.** Serum levels of liver transaminases (AST, ALT and ALP), albumin, triglyceride, cholesterol and glucose were assessed. Elevation of liver transaminases with reduction in serum albumin in PEX1-G844D homozygotes relative to littermate controls (N=3-7 per genotype). Hypotriglyceridemia and hypoglycemia were observed in PEX1-G844D mutants. Samples were assessed at age 3-9 weeks, 10-16 weeks, 17-23 weeks, 24-30 weeks and 31-37 weeks. The single data point of PEX1-G844D mutant at age 3-9 weeks was obtained by pooling serum from 3 different mice together given its small body size and limited amount of blood that could be drawn at early age. Unpaired student t-test. * $P<0.05$; ** $P<0.01$; *** $P<0.001$.
